# Temporal AI model predicts drivers of cell state trajectories across human aging

**DOI:** 10.64898/2026.03.30.715396

**Authors:** Javier Gόmez Ortega, Rangarajan D. Nadadur, Akira Kunitomi, Steven Kothen-Hill, Julian U. G. Wagner, Sarp D. Kurtoglu, Bumjoon Kim, Madigan M. Reid, Thomas Lu, Kaho Washizu, Lukas Zanders, Han Chen, Yujie Zhang, Sarah Ancheta, Sara Lichtarge, William A. Johnson, Chris Thompson, Dereck M. Phan, Alexis J. Combes, Andrew C. Yang, Neha Tadimeti, Stefanie Dimmeler, Shinya Yamanaka, Michael Alexanian, Christina V. Theodoris

## Abstract

Foundational AI models have recently shown promise for predicting the impact of perturbations on cell states. However, current models typically consider only one cell state at a time, limiting their ability to learn how cellular responses unfold over time, particularly across long trajectories such as diseases of aging. Here, we develop a temporal AI model, MaxToki, trained on nearly 1 trillion gene tokens including cell state trajectories across the human lifespan to generate cell states across long timelapses of human aging. MaxToki generalized to unseen trajectories through in-context learning and predicted novel age-modulating targets that were experimentally verified to influence age-related gene programs and functional decline in vivo. MaxToki represents a promising strategy for temporal modeling to accelerate the discovery of interventions for programming therapeutic cellular trajectories.

## Main Text

Mapping the gene networks disrupted in human disease enables the design of network-correcting therapies that restore disease-dependent networks back to the normal state (*1*, *2*). However, mapping these networks using traditional methods is commonly impeded by limited data, especially in rare diseases and diseases affecting clinically inaccessible tissues. To overcome this, we previously developed a transfer learning strategy for network biology where we first pretrained an artificial intelligence (AI) model, Geneformer, to gain a foundational understanding of gene network dynamics through observing gene network states across a broad range of human tissues in health and disease (initially ∼30 million in 2021, now >100 million) (*3*, *4*). We then designed an in silico perturbation strategy leveraging Geneformer to enable context-aware predictions in network biology. Geneformer drove biological insights including discovering a novel transcription factor in cardiomyocytes with zero-shot learning and predicting candidate therapeutic targets for cardiomyopathy that were experimentally verified to impact contractility in an induced pluripotent stem cell (iPSC) model of disease (*3*).

Recently, there have been multiple foundation models developed leveraging transfer learning to enable predictions in a diverse array of downstream tasks relevant to network dynamics (scBERT, tGPT, scGPT, scFoundation, GeneCompass, UCE, Nicheformer, scSimilarity, TranscriptFormer, STATE, and more) (*5–14*). Yet, a key limitation of Geneformer and related models is that they consider only a single cell state at a time, while in actuality cellular responses unfold over time, dynamically progressing across cell state trajectories in development, aging, and disease. Understanding how gene network perturbations affect not just the current cell state but that cell’s trajectory over time would enable predicting interventions to induce desired cell state transitions for both cellular engineering and for programming lasting therapeutic cellular trajectories.

Particularly for aging, which progresses across very long timelapses, AI forecasting of cellular responses would allow large-scale in silico screening of candidate interventions to slow age-dependent decline in tissues affected by age-related disease, nominating promising targets in advance of costly long-term clinical trials. Diseases of aging, including heart disease, dementia, and cancer, are the number one cause of morbidity and mortality worldwide and are only increasing in prevalence with the aging population (*15*). While sampling the same individual over time in disease-relevant tissues is generally not possible, assembling a population-based corpus of gene network changes over time across many individuals may enable AI models to learn the shared trajectories of age-dependent decline where intervention would also be most broadly effective across individuals.

Here, we develop a temporal AI model, MaxToki, for generating past, intervening, and future cell states across dynamic trajectories and in silico predicting interventions to induce desired cell state transitions. We designed a two-stage strategy where MaxToki was first trained with ∼175 million single-cell transcriptomes to generate cell states and then trained with ∼100 million cell state trajectories to generate cells and predict timelapses between cell states across human aging from birth to the tenth decade of life. We implemented advances in GPU acceleration to enable the modeling of multiple cells in time series, training a 217 million and 1 billion parameter model with nearly 1 trillion gene tokens across the two-stage training. We formulated a prompting strategy to query the trained model and found that MaxToki learned to predict timelapses needed to induce cell states from both held-out ages and held-out cell types as well as to generate cell states that matched the prompted age and cell type. Interpretability analysis revealed the model learned in a completely self-supervised manner to pay higher attention to transcription factors, which are critical drivers of cell state trajectories, and that the model attended to both the context and query in the prompt to optimize its responses. The model inferred age acceleration in diseases of aging never seen during training, including pulmonary fibrosis and Alzheimer dementia, and predicted candidate age-promoting vs. rejuvenating perturbations in cardiac cell types relevant to age-related cardiovascular decline. Top novel pro-aging perturbations were experimentally validated to cause age-related gene network dysregulation in cardiomyocytes and functional decline both in vitro and in vivo. Overall, MaxToki represents a temporal AI model for predicting the drivers of cell state progression over time, providing a generalizable framework to decode and control dynamic cellular trajectories.

### MaxToki architecture and pretraining

MaxToki is a temporal AI model designed to generate past, intervening, and future cell states across dynamic trajectories (Fig. 1A). The model undergoes a two-stage training, first learning to generate single cell transcriptomes and then expanding the input size to model multiple cell states along a context-specific trajectory. By learning how gene network states direct future cell state transitions, the model can be prompted to generate cells along the continuum or predict how perturbations would impact the trajectory.

**Fig. 1.**
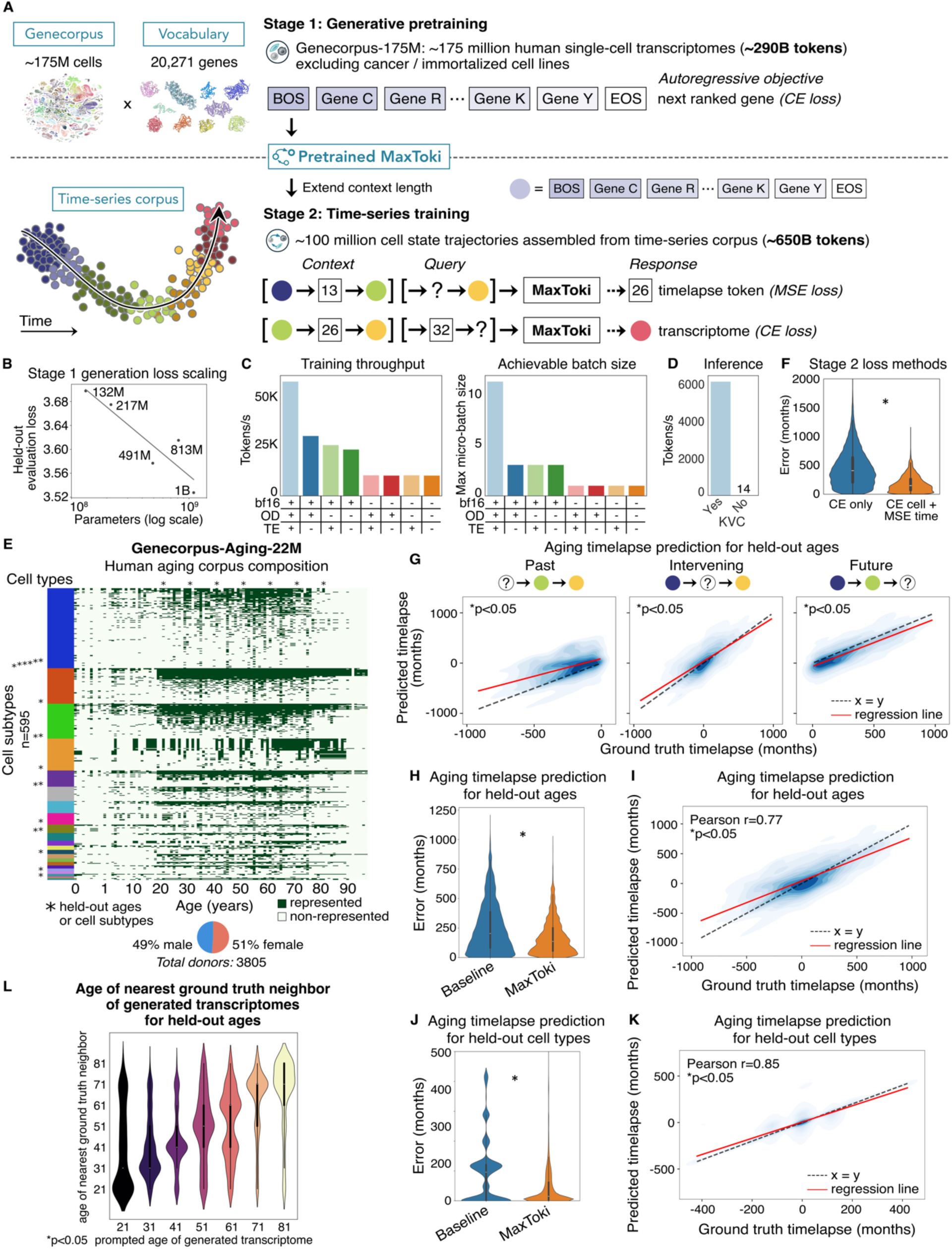
MaxToki learned to generate cell states and predict the timelapse needed to induce cell states across long trajectories of human aging. (A) Two-stage training strategy of MaxToki. The model is trained in total on nearly 1 trillion gene tokens (non-trivial tokens, excluding non-expressed genes). MSE=mean-squared error, CE=cross-entropy. (B) Stage 1 generation loss scaled with increasing model parameters. (C) Implementing advances in GPU acceleration including mixed precision training with bf16 (vs. fp32), optimizing model dimensions (OD), and Transformer Engine (TE) improved model throughput by 5x and allowed 4x larger batch sizes due to reduced memory usage. (D) Implementing key-value caching (KVC) led to >400x improved generation speed. (E) Composition of population-based human aging corpus of single-cell transcriptomes over the human lifespan from birth to the tenth decade of life across ∼3800 individuals and ∼600 cell types. Asterisks indicate cell types and ages held-out during training that are used for evaluation in (F-L). (F) Stage 2 timelapse prediction error for held-out ages when using CE loss for both tasks vs. using CE for cell generation and MSE for timelapse prediction with continuous numerical tokenization (n=50,092). (G) Aging timelapse prediction for held-out ages where the query cell represents a past *(left)*, intervening *(middle)*, or future *(right)* cell state along the trajectory with respect to the context cells within the prompt (n=44,918) (H) Aging timelapse prediction for held-out ages compared to baseline approach of predicting timelapses by assuming the query cell was the most common age of that cell type and sex (n=44,918). (I) Correlation of predicted vs. ground truth aging timelapses for held-out ages (n=44,918). (J) Aging timelapse prediction for held-out cell types compared to baseline approach as described in (H) (n=46,000). (K) Correlation of predicted vs. ground truth aging timelapses for held-out cell types (n=46,000). (L) Age of the nearest ground truth neighbor of generated transcriptomes prompted for held-out ages, as evaluated in an external model’s embedding space (n=4988). (F,H,J): *p<0.05, Wilcoxon rank sums. (G,I,K): *p<0.05, Pearson.

To accomplish this, we first assembled Genecorpus-175M, a large-scale pretraining corpus composed of ∼175 million single-cell transcriptomes from a broad range of human tissues in health and disease from publicly available data (Table S1). We balanced the data such that no tissue composed more than 25% of the data and deduplicated the cells by DOI. Cells with high mutational burdens like malignant cells and immortalized cell lines were excluded as they are known to contain gain of function variants that alter gene functions from what the model would observe in other cells with low mutational burdens.

Each cell transcriptome was presented to the model as a rank value encoding, a non-parametric representation of the transcriptome where genes are ranked by their relative expression in a given cell after scaling by their expression across the entire pretraining corpus. This method deprioritizes ubiquitous highly expressed housekeeping genes while prioritizing genes such as transcription factors that may be generally lowly expressed but have a high dynamic range across distinct cell states. Furthermore, the rank-based approach may be more robust against technical artifacts that may systematically bias the absolute transcript counts value whereas the overall relative ranking of genes within each cell remains more stable.

We then pretrained MaxToki, a transformer decoder model, with an autoregressive objective to generate the next ranked gene across each single-cell transcriptome. We first partially trained a series of models of different parameter sizes to define the scaling laws of generative transcriptome modeling. We found that models with a larger number of parameters scaled as a power law in their performance on the autoregressive objective of generating single-cell transcriptomes. To balance performance with compute budget constraints, we fully pretrained two variants: a 217 million (M) parameter variant pretrained in full precision and a 1 billion (B) parameter variant pretrained with mixed precision, both with an initial context size of 4096.

To enable efficient usage of the 1B parameter variant, we optimized training and inference using components of the NVIDIA BioNeMo stack, built on NeMo, Megatron-LM, and Transformer Engine. To enable FlashAttention-2, we modified model parameter shapes so that feed-forward hidden dimensions were compatible with attention head partitioning by ensuring the feed-forward hidden dimensions evenly divided the number of attention heads. We then activated FlashAttention-2 through the integrated stack, leveraging fused kernels across NeMo, Megatron-LM, and Transformer Engine. Transformer Engine reduced memory footprint and improved arithmetic throughput through kernel fusion and mixed-precision execution. Together, these changes yielded approximately a 5x increase in training throughput and a 4x increase in achievable micro-batch size on H100 80GB GPUs (Fig. 1C).

For inference, we adopted the Megatron-Core inference engine using the DynamicInferenceContext abstraction. This object manages key–value cache allocation and reuse across decoding steps and supports sequence packing for efficient utilization of compute and memory. These optimizations resulted in a >400x improvement in autoregressive generation speed relative to the baseline implementation (Fig. 1D).

### Temporal training and prompting strategy

After pretraining the model to generate individual single-cell transcriptomes, we then extended the context length to 16,384 with RoPE scaling to accommodate an input of multiple single-cell transcriptomes along a cell state trajectory. RoPE scaling modifies the Rotary Positional Embeddings of the model by reducing the frequency of rotation to interpolate more tokens into the existing positional framework (*16*). Within the extended input size, the model is presented with a series of multiple cells across a cell state trajectory along with the time elapsed between each state. Special tokens are added to indicate the start and end of each given cell state as well as the start and end of the query posed to the model during training and inference.

To train the model to understand how cell states progress along context-specific trajectories, we designed a temporal prompting strategy where the model first learns from an initial trajectory for a given context (e.g. given cell type and sex) and is then tasked with either 1) predicting the time needed to elapse to lead to a query cell or 2) generating the cell that would arise after a query timelapse along the prompted trajectory (Fig. 1A). For the task of cell state generation (task 2), the model is trained to generate the next ranked gene with a cross-entropy loss function, analogous to the pretraining objective. However, for the task of predicting the timelapse needed to generate a query cell (task 1), we employ a form of continuous numerical tokenization (*17*) with a mean-squared error loss function that rewards predictions closer to the target value so that the model can learn that timelapse tokens fall along a numerical continuum. The model learns from these two cross-informative tasks to understand how gene network states progress along the context-specific trajectory, inferring the context from the initial prompt without explicit supervision.

### Assembly of population aging corpus across every decade of life

We first sought to apply MaxToki to learn common mechanisms of gene network dysregulation that occur over the long timelapses of human aging. Age-related decline involves progressive changes in cell states that unfold over time, but in most tissues, we are unable to serially measure gene expression profiles within the same individual over the course of aging. To address this, we took the approach of assembling a population-based aging corpus from publicly available data comprising ∼22M cells across ∼600 human cell types from ∼3800 donors representing every decade of life from birth to 90+ years (Fig. 1E). The donors were balanced by sex and annotated in the original studies as controls with no known diseases.

From this data, we constructed ∼100M simulated aging trajectories across individuals stratified by cell type and sex. Each example was composed of an initial trajectory of 2-3 cells and either a query cell or query timelapse, where the model was prompted to predict the time elapsed between states or generate the cell that would arise after the given timelapse, respectively (tasks 1 and 2 described above). The timelapses in the example trajectories included both positive and negative time to enable prompting the model to generate past and intervening cell states in addition to future cell states along the trajectories. In addition, we held-out a portion of cell types and ages from the corpus to later confirm generalizability of the trained model.

### MaxToki predicted timelapses between aging cell states for held-out ages and cell types

We utilized the population-based aging corpus to train MaxToki to generate cell states and the timelapses between them across the trajectory of human aging (Fig. S1A). We found that the method for continuous numerical tokenization significantly improved the ability of the model to learn to predict the timelapses between cell states compared to using standard cross-entropy loss for both tasks (Fig. 1F). By learning that timelapses occur along a numerical continuum, the model was able to predict the timelapse between cells within seen trajectories even if the order of cell states was shuffled, changing the correct response for the relative timelapse between cell states (Fig. S1B-C). In contrast, a linear model approach (SGDRegressor) was not able to sufficiently improve predictions compared to the baseline approach of predicting timelapses by assuming the query cell was the most common age of that cell type and sex (median error: baseline: 180, SGDRegressor: 178, MaxToki: 87) (Fig. S1B-C).

We then evaluated the model’s ability to predict the timelapses between aging cell states for query cells from held-out ages (and therefore also held-out donors). We found that the model was able to predict the timelapse to query cells that were either past, intervening, or future cell states within the trajectory provided in the prompt’s context with significant correlation between the predicted and ground truth timelapses (Fig. 1G). Overall, the correlation between ground truth and predicted timelapses was 0.77, significantly improving the error compared to a linear model (SGDRegressor) as well as the baseline approach of predicting timelapses by assuming the query cell was the most common age of that cell type and sex (Fig. 1H-I, Fig. S1D-E).

Similarly, the model was able to accurately predict the timelapse between cell states of held-out cell types based on in-context learning of the unseen trajectory, significantly improving predictions compared to the baseline approach with correlation to ground truth of 0.85 (Fig. 1J-K).

### MaxToki generated cell states across aging consistent with the prompted timelapse

Next, we evaluated the cell transcriptomes generated by the model in response to prompts of a given timelapse along the context-specific trajectory. The model was prompted to generate the rank value encoding representation of cells of held-out ages that the model had not seen during training. When embedding these generated cells along with corresponding ground truth cells in an external model’s embedding space, we found that indeed the nearest ground truth neighbor to the generated cells matched the expected age prompted by the timelapse (Fig. 1L).

Although the cell type context is not explicitly labeled in the prompt, the model must infer the correct cell type context to generate from the initial trajectory in the prompt. Indeed, the generated cells prompted to represent cells of held-out ages also matched the cell type context in the prompted trajectory. When an external model cell type classifier was applied to both the ground truth and generated cells, there was an 82% concordance of the cell type annotated by the model for each group (Fig. 2A). Consistently, generated cells were co-embedded with ground truth cells of the same cell type within the external model’s embedding space (Fig. 2B).

**Fig. 2.**
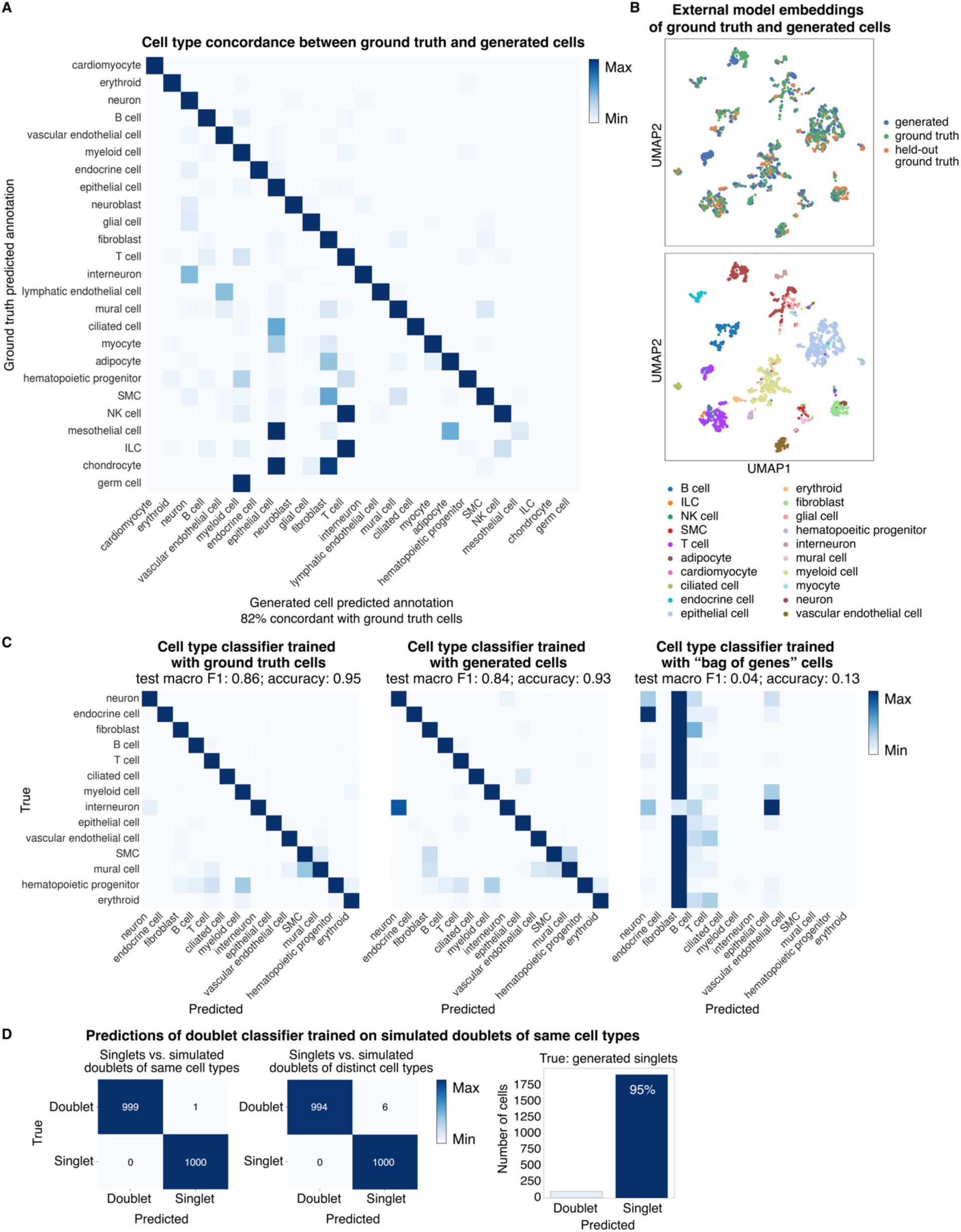
Generated cell states were consistent with the prompted cell context. (**A**) Cells generated in response to a prompted contextual trajectory were annotated by an external cell type classifier model with 82% concordance with annotations of ground truth cells of that cell type (n=87,083). (**B**) Generated and ground-truth cells of the same cell type were co-embedded in an external model’s embedding space. (**C**) Newly initialized cell type classifiers were trained with either ground truth cells, generated cells, or “bag of genes” cells where gene rank order in ground truth cells was shuffled (n=4000). Models trained with either ground truth or generated cells were able to accurately classify the cell type of held-out ground truth cells, indicating the generated cells contained key features for learning cell identity. In contrast, models trained on “bag of genes” cells were unable to annotate held-out ground truth cells, indicating the binary presence of cell type markers without correctly representing their relative expression levels in each cell type was insufficient to teach the model to classify held-out cells. (**D**) Generated cells were predominantly classified as singlets by a classifier trained to distinguish single cells from simulated doublets of the same cell type, indicating the generated cells represented single-cell resolution (n=2000).

An even stricter test of the fidelity of the generated cells would be to train an external model on either ground truth or generated cells and then test its ability to classify the cell type of held-out ground truth cells. Only if the informational content of the generated cells faithfully captured key features of ground truth cells would a model trained solely on generated cells be able to annotate unseen ground truth transcriptomes. Indeed, a cell type classifier trained with generated cells achieved a near equivalent accuracy on annotating held-out ground truth cells as compared to a classifier trained with true cells (Fig. 2C). Furthermore, a classifier trained on “bag of genes” cells, in which the order of genes within the ground truth rank value encoding was shuffled, was unable to classify held-out ground truth cells, indicating that the generated cells captured not only the presence of marker genes but also critical components of the true rank value order of the transcriptomes.

Finally, we tested the resolution of the generated cells. One common pitfall of generative models is learning to generate an average output to minimize the loss function. To evaluate this, we first trained a classifier to distinguish singlets vs. simulated doublets of the same cell types from ground truth cells (individual cells vs. merged cells from the same cell type). We confirmed that the trained classifier could accurately distinguish singlets vs. simulated doublets from held-out ground truth cells of the same cell type or distinct cell types. We then applied this classifier to the generated cells and found that the vast majority of cells were classified as singlets, indicating that the model produces rank value encodings consistent with an individual single-cell rather than merging multiple cells into an average encoding (Fig. 2D).

### MaxToki generated cell states capturing non-monotonic gene activation within held-out inflection points in short timelapse trajectories

Having confirmed the model could predict timelapses and generate transcriptomes across long timelapses of aging, we then tested the model’s ability to generate cells within held-out inflection points in short timelapse trajectories. We trained MaxToki using pseudotime trajectories of partial reprogramming of primary adult human cytotoxic T lymphocytes, dermal fibroblasts, and aortic endothelial cells. The partial reprogramming with OCT4, SOX2, KLF4, and c-MYC (OSKM) represents a rejuvenating perturbation that partially reprograms cells without fully inducing pluripotency. Three stages of reprogramming were defined as stage 1: positive for somatic identity marker and negative for pluripotency marker, stage 2: negative for both somatic identity marker and pluripotency marker, and stage 3: negative for somatic identity marker and positive for pluripotency marker.

We held out pseudotime intervals corresponding to the inflection points between these stages and tested whether the model could predict the pseudo-timelapses needed to induce the inflection point and generate cell states within these held-out intervals (Fig. 3A). We found that indeed MaxToki was able to predict the pseudo-timelapse needed to induce cell states within these held-out intervals along the trajectory, significantly improving predictions compared to baseline and strongly correlating between ground truth and predicted pseudo-timelapses (Fig. 3B-C, S1F). We then prompted the model to generate cell states along the trajectory that would fall into the held-out intervals in the T cell trajectory. We identified the genes that were predicted by the model to be most overexpressed or repressed within the held-out intervals defined as inflection points to stage 2 and 3 by comparing the gene ranks in the cells generated within those held-out intervals vs. all other cells across the trajectory. We found that these top-most overexpressed or repressed genes were consistent with a non-monotonic pattern of increasing or decreasing expression, respectively, within the inflection points in the ground truth cells (Fig. 3D-F). Also notable was that these top differentially expressed genes within the generated cells included genes that appeared to have dynamic changes in ground truth expression at each of the three stage inflection points, suggesting that they may correspond to gene programs associated with cell fate transitions.

**Fig. 3.**
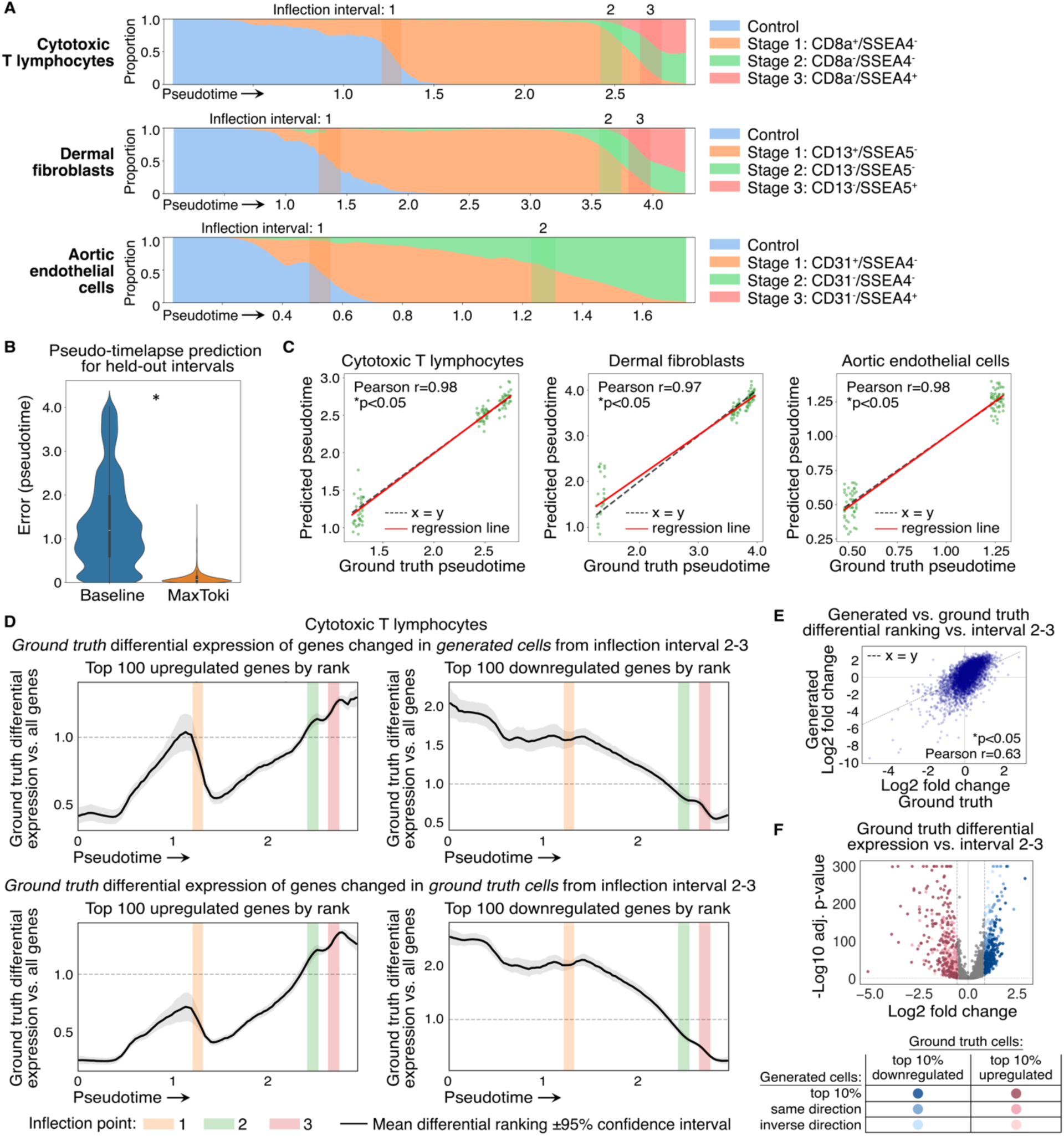
MaxToki generated cell states capturing non-monotic gene activation in held-out inflection intervals within short timelapse trajectories. (**A**) Diagram of held-out inflection intervals across partial reprogramming pseudotime trajectories in the indicated cell types. The plots indicate the proportion of cells at each point in pseudotime representing cells annotated by cell markers for each stage of partial reprogramming. While the proportion of cells in each stage may appear monotonic, gene expression levels may change non-monotonically within the held-out intervals, as indicated in (D). (**B**) Pseudo-timelapse prediction for held-out intervals compared to baseline of predicting most common pseudo-timelapse for that cell type. *p<0.05, Wilcoxon rank sums (n=25,000). (**C**) Correlation of predicted vs. ground truth pseudotime for each cell type in held-out intervals (n=100). (**D**) Ground truth differential expression of top 100 upregulated or downregulated genes by rank within generated *(top)* or ground truth *(bottom)* cells from inflection intervals 2-3 in cytotoxic T lymphocytes. (**E**) Generated vs. ground truth differential ranking of genes vs. interval 2-3 in cytotoxic T lymphocytes (n=4463). (**F**) Ground truth differential expression vs. interval 2-3 indicating overlap with differential ranking in generated cells (Wilcoxon rank sums with Benjamini-Hochberg (BH) correction, n=4463). (C,E): *p<0.05, Pearson.

### Interpretability analysis revealed specialization of attention heads across the context, timelapse, and query features and higher attention to central regulators

We then performed an interpretability analysis to determine how MaxToki was utilizing the information within the prompt to accurately predict the timelapses between cell states along the trajectory. For this analysis we focused on the model trained with the long timelapse trajectories across aging and analyzed its predictions of timelapses for held-out ages and cell types. We first performed ablation studies to identify the input features that were most critical to the model predictions. We found that when the cells within the prompt were presented as “bag of genes” cells (maintaining the presence of genes expressed within each cell but shuffling their order within the rank value encoding), this significantly damaged the ability of the model to predict cell state timelapses for held-out cell types (Fig. 4A). Furthermore, both the context and query were equally important in achieving accurate timelapse predictions, as masking either component significantly damaged predictions (Fig. 4B).

**Fig. 4.**
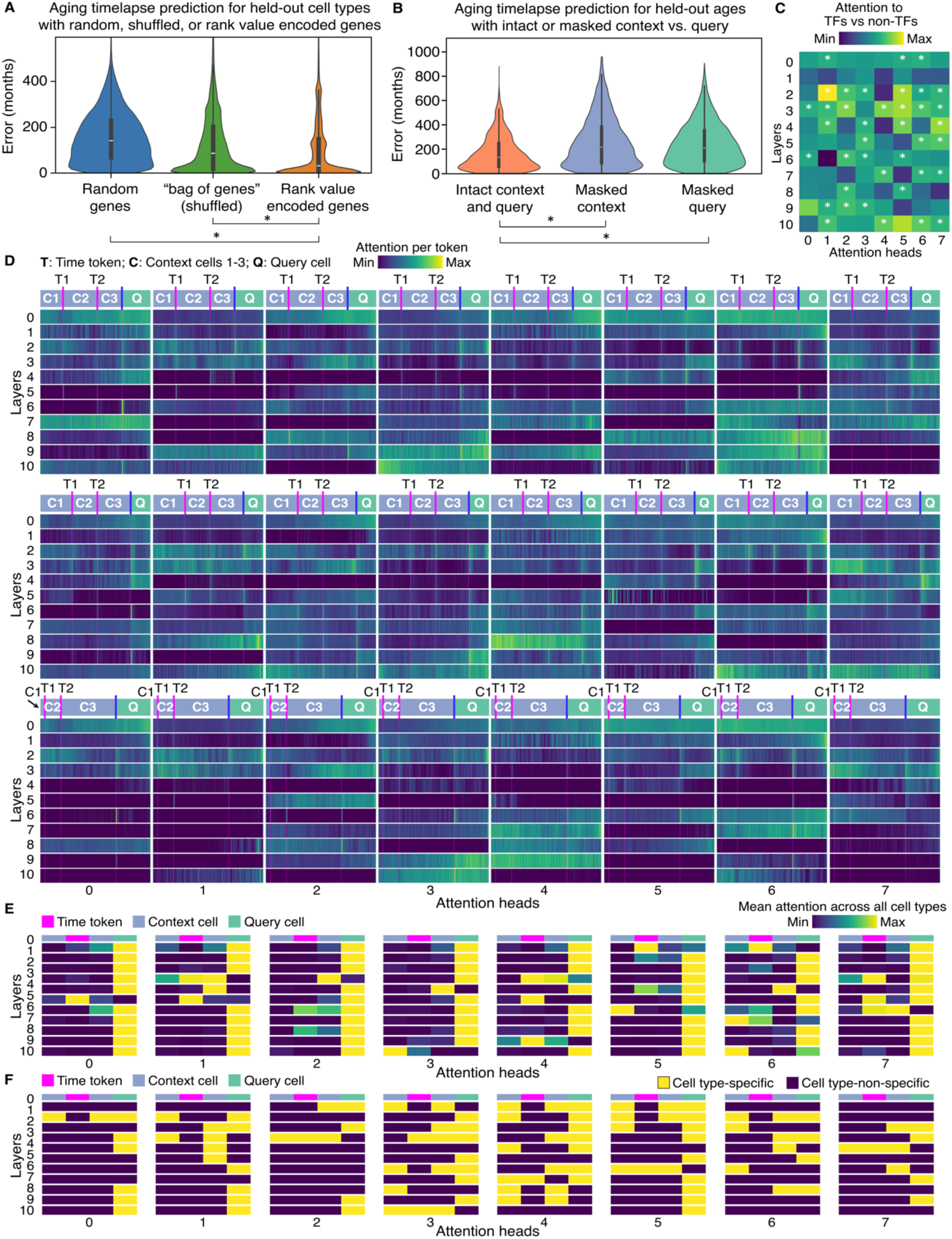
Interpretability analysis revealed context-specific specialization of attention heads and higher attention to central regulators. (**A**) Aging timelapse prediction for held-out cell types was significantly damaged by shuffling gene order within the rank value encoding, indicating that relative gene expression levels in each cell state were essential for in-context learning of unseen trajectories (n=50,000). (**B**) Aging timelapse prediction for held-out ages was equivalently damaged by masking the context or the query, indicating the model uses information from both components of the prompt for inferring the correct timelapses and does not only rely on the query cell (n=2619). (**C**) Multiple attention heads learned in a completely self-supervised manner to pay significantly higher attention to transcription factors (TFs) (which are known key regulators of cell state trajectories), compared to other genes, despite no prior information about gene function. *p<0.05, Wilcoxon rank sums with BH correction, n=1000. (**D**) Three example prompts of an initial contextual trajectory composed of three cell states and the timelapses between them followed by a query cell state. Attention weights across the prompt sequence length are shown for each attention head in each layer of the model. Each attention head pays attention to variable regions of the sequence, with some attention heads exhibiting context-specific attention that differs between the three example prompts. (**E**) Mean attention across all cell types to each key component of prompts composed of two context cells and the timelapse between them followed by a query cell state (n=1000). (**F**) Attention was analyzed for prompts across ∼60 random cell types to quantify attention heads that paid significantly higher attention to one cell type compared to the others within each of the indicated prompt components (context cell 1, time token, context cell 2, and query cell). *p<0.05, Wilcoxon rank sums with BH correction, n=1000 (n>10 for each cell type). (A-B): *p<0.05, Wilcoxon rank sums. (A-F): interpretability analysis was performed with the 217M parameter model.

We then analyzed the model attention weights to uncover how each of the attention heads in each layer jointly contributed to the model predictions. Firstly, we found that about half of the attention heads paid significantly higher attention to transcription factors compared to other genes (Fig. 4C). This indicated that the model had learned in a completely self-supervised manner without prior information about gene function to pay higher attention to these known important regulators of cell state transitions.

When we mapped the attention weights for each position along the prompt including three context cells and the timelapses between them as well as the query cell, we noted that each attention head learned to pay attention to a different region of the sequence to optimize the model predictions (Fig. 4D-E). Some attention heads paid higher attention to the context cells, some paid higher attention to the timelapse tokens, and others paid higher attention to the query cell. The model appeared to recognize that these features were coherent units of information, as certain heads that paid higher attention to a given feature, such as the query cell, shifted their attention to different regions of the sequence dependent on the length occupied by that feature in each specific example. Furthermore, many heads paid high attention to the boundaries between input features, again shifting attention to the position of the boundaries in each example prompt.

Interestingly, some attention heads appeared to be context-specific, variably activating depending on the trajectory context. For instance, Layer 8, Head 4 paid low attention to the context cells in the first example prompt shown in Fig. 4D but high attention in the second example and intermediate attention in third. On the other hand, Layer 4, Head 7 paid high attention to the query cell in the first and second examples and lower attention in the third. In fact, when we tested prompts composed of cells from each of about 60 example cell types, we found that most heads paid attention to at least one input feature significantly more in at least one cell type (Fig. 4F).

Overall, the interpretability analysis found that the model paid attention to a complex array of features within the prompt including the trajectory context, query cell, gene rank value order, and central regulators to optimize its predictions of timelapses to held-out cell states.

### MaxToki inferred age acceleration of cell states in age-related diseases

Given the model could predict the timelapses between held-out cell states for the trajectory of normal aging, we then tested the model’s interpretation of disease states relevant to age-related dysfunction. We perform this analysis by presenting the model with a context trajectory of normal cells followed by a query cell that is either from a donor affected by a given disease or from an age-matched unaffected donor. We then test whether the inferred timelapse to the disease cell state is longer or shorter than the true timelapse based on the donor’s chronological age, compared to the timelapse inferred to an age-matched control cell state.

We first tested whether the model would interpret that mucosal epithelial cells in the lung of donors exposed to heavy smoking were younger or older than non-smoking age-matched controls (*18*). Indeed, we found that the model interpreted about a five year age acceleration in the lung cells from donors exposed to heavy smoking compared to age-matched controls. This inferred age acceleration in the lung cells of heavy smokers is consistent with prior reports that aging signatures, lung dysfunction over time, and telomere length are all impacted by smoking status (*19*) (Fig. 5A-B).

**Fig. 5.**
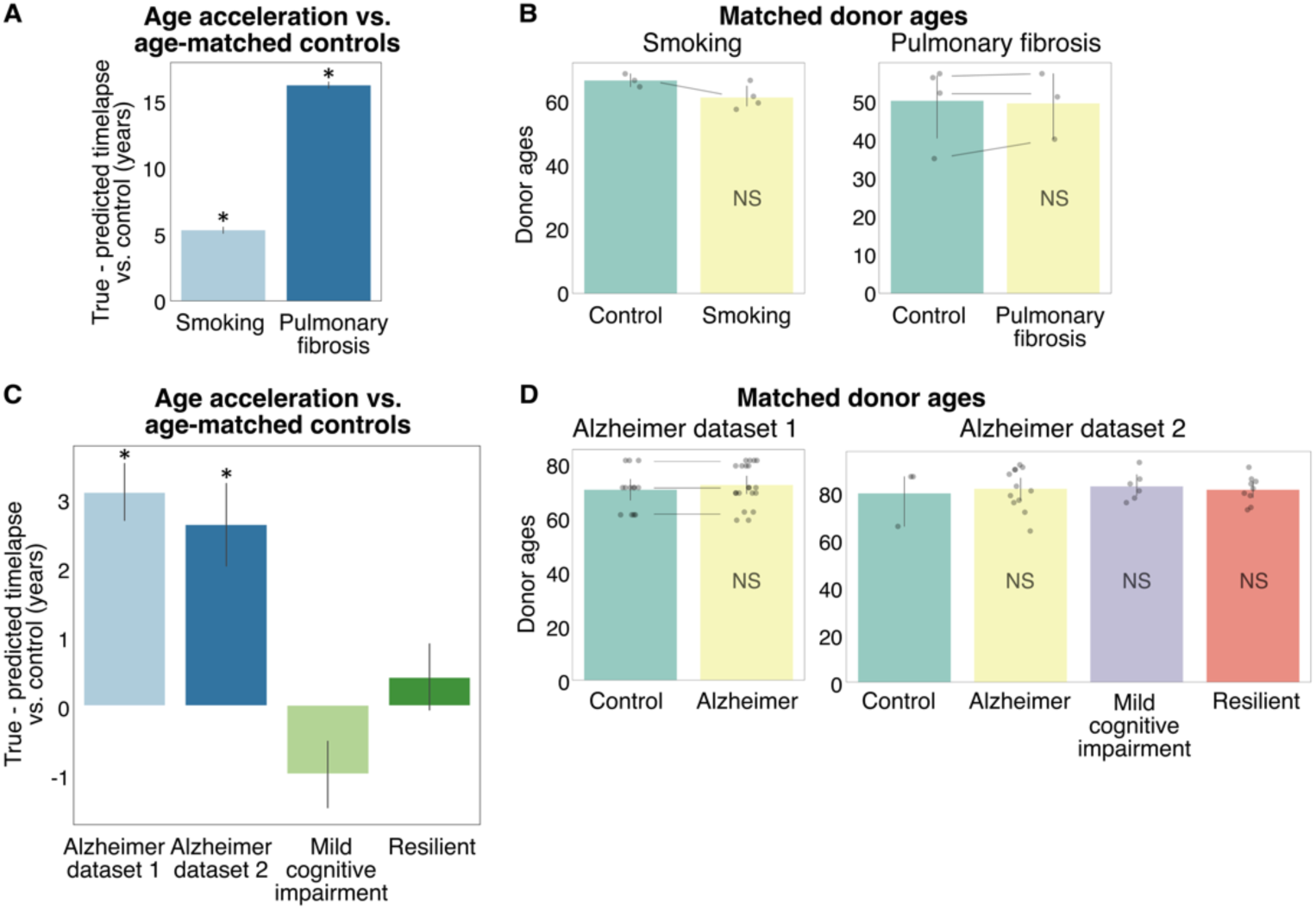
MaxToki inferred age acceleration of cell states in age-related conditions. (**A**) Inferred age acceleration of disease cell states vs. age-matched controls in donors exposed to heavy smoking (lung mucous secretory epithelial cells) or affected by pulmonary fibrosis (lung fibroblasts) (n=5000). (**B**) Matched donor ages for disease vs. control query cells used for analysis shown in (A). Lines indicate comparison subgroups. (**C**) Inferred age acceleration of disease cell states vs. age-matched controls in patients affected by Alzheimer dementia, mild cognitive impairment, or resilient patients who have the same neuropathology as Alzheimer patients with no cognitive impairment (dataset 1: microglia, n=5000; dataset 2: homeostatic microglia, n=2507). (**D**) Matched donor ages for disease vs. control query cells used for analysis shown in (B). Lines indicate comparison subgroups. (A-D): *p<0.05, Wilcoxon rank sums. NS=non-significant.

We then tested whether the model would infer that lung fibroblasts from patients with pulmonary fibrosis would be younger or older than age-matched controls (*20*). Pulmonary fibrosis is a known age-related disease that involves telomere attrition, cellular senescence, and decreased regenerative capacity of the lungs (*21*). Compared to age-matched controls, the model inferred a 15 year age acceleration in the lung fibroblasts from pulmonary fibrosis, consistent with the known aging effect of the disease (Fig. 5A-B).

Finally, we tested whether the model would infer that microglia from patients with Alzheimer disease were younger or older than age-matched controls. Microglia are known to exhibit age-related changes in Alzheimer disease and are a therapeutic target of interest for mitigating disease progression (*22*, *23*). We first tested microglia from Alzheimer patients vs. age-matched controls from the Mount Sinai NIH Neurobiobank (*24*) and found that microglia from Alzheimer patients were inferred to have a three year age acceleration (Fig. 5C-D).

We then replicated this analysis in an external cohort of Alzheimer patients from the Duke and Johns Hopkins Alzheimer Disease Research Centers. This second cohort included fine annotations of microglia states so we tested each state as a separate context. Again, we found that the model interpreted similar age acceleration in affected patients specifically in microglia annotated as the homeostatic subtype (Fig. 5C-D). The second cohort also included age-matched patients with mild cognitive impairment or Alzheimer resilience, which is characterized by exhibiting the same level of neuropathological changes with no cognitive impairment. Interestingly, homeostatic microglia from those patients who showed only mild cognitive impairment or Alzheimer resilience were not interpreted by the model to exhibit age acceleration compared to age-matched controls, suggesting these patients were protected from the disease-related age acceleration in this microglial subtype (Fig. 5C-D).

### In silico prediction and experimental validation of pro-aging drivers in cardiac cell types

The ultimate goal of developing a model to understand the progression of cell states along trajectories is to enable predictions of candidate perturbations to induce desired trajectory shifts. We therefore designed an in silico perturbation strategy to predict age-promoting vs. rejuvenating targets by applying perturbations to the query cell within a given prompt and testing whether the model inferred that the timelapse needed to induce the perturbed cell would be shorter or longer than the unperturbed trajectory.

We first applied this approach to predict age-promoting vs. rejuvenating targets in cardiac fibroblasts, which are a key contributor to age-related cardiovascular decline via pro-fibrotic remodeling and dysfunctional repair pathways (*25*). We found that the predicted impact of in silico repression significantly inversely correlated with the slope of expression change across human aging (Fig. 6A-B). Interestingly, one of the top predicted rejuvenating perturbations was repression of GSN, a gene whose inhibition was previously validated to improve contractility of cardiac microtissues in a model of dilated cardiomyopathy (*3*) and whose expression increases over the course of human aging. Overall, gene set enrichment of in silico predicted targets in cardiac fibroblasts included known aging pathways such as the mTOR and AGE-RAGE signaling (*26*, *27*), pathways relevant to dysfunctional fibroblast activation such as TGFbeta stimulus (*28*) and response to stress, and emerging pathways contributing to age-related cardiovascular decline such as the Relaxin (*29*) and glycosaminoglycan binding (*30*) (Fig. S2A, Table S2). Of particular potential therapeutic interest was the gene set involved with extracellular vesicles in the crosstalk of cardiac cells as these targets may serve to rejuvenate not only the fibroblasts but also surrounding cells within the cardiac tissue.

**Fig. 6.**
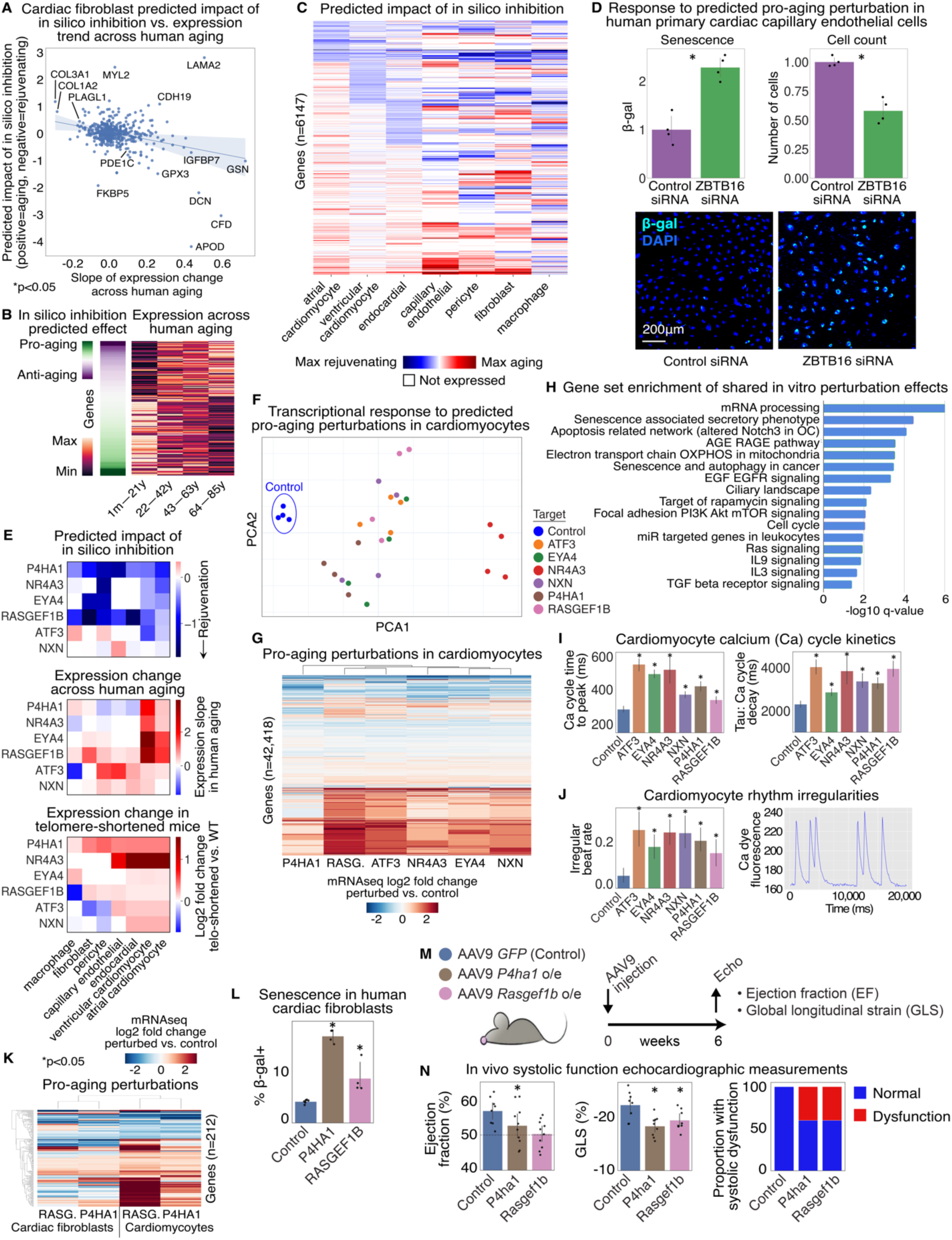
In silico perturbation predicted pro-aging perturbations that were experimentally verified in cardiac cell types. (**A**) Predicted impact of in silico inhibition vs. slope of expression change across human aging for genes in cardiac fibroblasts. (**B**) Concordant genes whose inhibition was predicted to be rejuvenating were upregulated across aging, and vice versa. (**C**) Predicted impact of in silico inhibition of genes across the indicated cardiac cell types. (**D**) Beta-gal staining and cell counts quantifying senescence of primary human cardiac capillary endothelial cells in response to predicted pro-aging perturbation of ZBTB16 inhibition. *p<0.05, Wilcoxon rank sums, n=4. (**E**) Predicted impact of in silico perturbation and expression change across human aging and in telomere-shortened mice in each cardiac cell type for top predicted cardiomyocyte age-modulating targets. (**F**) PCA plot and (**G**) differential gene expression heatmap of transcriptional response (measured by bulk RNA sequencing) to predicted pro-aging perturbations (AAV overexpression of indicated genes vs. GFP) in human iPSC-derived cardiomyocytes. p<0.05, Wald test with BH correction, n=4. (**H**) Gene set enrichment of genes differentially expressed in response to all predicted pro-aging perturbations in human iPSC-derived cardiomyocytes (hypergeometric test with g:Set Counts and Sizes (g:SCS) correction). (**I**) Slowed calcium cycle kinetics and (**J**) rhythm irregularities in response to predicted pro-aging perturbations (AAV overexpression of indicated genes vs. GFP) in human iPSC-derived cardiomyocytes. *p<0.05, Wilcoxon rank sums with BH correction. Rhythm n=10, time to peak n=180, 111, 139, 37, 237, 150, 191 (bars left to right), decay n=169, 30, 86, 11, 213, 105, 172 (bars left to right). (**K**) Differential gene expression heatmap of transcriptional response (measured by bulk RNA sequencing) to predicted pro-aging perturbations (AAV overexpression of indicated genes vs. GFP) in human iPSC-derived cardiomyocytes or human primary cardiac fibroblasts where dysregulation of the given gene was statistically significant across all four conditions. Statistical significance of differential expression: p<0.05, Wald test with BH correction, n=4. Statistical significance of overlap of shared dysregulation between the four conditions: p<0.05, one-sided binomial test. RASG.=RASGEF1B. (**L**) Beta-gal staining quantifying senescence of primary human cardiac fibroblasts in response to predicted pro-aging perturbations (AAV overexpression of indicated genes). *p<0.05, Wilcoxon rank sums with BH correction, empty n=6, P4HA1 and RASGEF1B n=4. (**M**) Schematic of in vivo validation experiment. o/e=overexpression. (**N**) In vivo systolic function echocardiographic measurements at 6 weeks post-AAV9 injection. EF: GFP n=8, P4ha1 n=10, Rasgef1b n=10; GLS: GFP n=8, P4ha1 n=9, Rasgef1b n=8. *p<0.05 one-way ANOVA with multiple hypothesis correction.

We then expanded our analysis to perform in silico perturbation screens in each of the major cardiac cell types to identify age-promoting vs. rejuvenating factors (Fig. 6C, Table S3). Clustering by perturbation effect across cell types revealed certain gene sets were predicted to be predominantly influential in a single cell type whereas others had a consistent predicted pro-aging or rejuvenating effect across all cell types. Predicted pro-aging perturbations in capillary endothelial cells included repression of ZBTB16, whose endothelial-specific knockout in mice was recently reported to cause premature aging, diastolic dysfunction, and secretion of pro-fibrotic and inflammatory factors (*31*). Consistently, when we tested its repression in human primary cardiac capillary endothelial cells, we found the cells experienced decreased cell counts and increased senescence, a hallmark of age-related dysfunction (Fig. 6D).

We next sought to experimentally test the top novel predicted targets in ventricular cardiomyocytes, as these cells are primary mediators of cardiac function and are particularly vulnerable to age-related decline. We selected five novel pro-aging perturbations along with a positive control, ATF3, whose activation in young mice was previously reported to cause age-related decline in cardiac function (*32*). In silico repression of each of the five targets was predicted to be rejuvenating in ventricular cardiomyocytes, correlating with their increased expression in ventricular cardiomyocytes across human aging and in telomere-shortened mice, which exhibit accelerated aging (Fig. 6E).

When we tested overexpression of the pro-aging targets in iPSC-derived cardiomyocytes, we found that all targets shifted the cell states away from the control-treated cells (GFP overexpression) and towards the positive control pro-aging perturbation of ATF3 (Fig. 6F, S2B-F, Table S4). The downstream transcriptional impact of the perturbations included a great portion of shared effects, with many genes being activated or repressed consistently across all six perturbations (Fig. 6G). Of the perturbations tested, overexpression of RASGEF1B clustered most closely with overexpression of the positive control ATF3, whereas P4HA1 was the most distinct in its transcriptional effects, as determined by hierarchical clustering.

Downstream differentially expressed genes shared across all predicted pro-aging perturbations were enriched for known aging pathways including mTOR and AGE-RAGE signaling (*26*, *27*), inflammatory pathways such as interleukin 3 and 9 signaling, genes involved in mitochondrial function, and the senescence-associated secretory phenotype, a program marked by secretion of pro-inflammatory factors that drive tissue dysfunction (*33*) (Fig. 6H, Table S5).

Furthermore, the transcriptional changes were associated with functional defects in the calcium cycle kinetics of the cardiomyocytes. Predicted pro-aging perturbations significantly prolonged the time to peak and decay kinetics of the calcium transient (Fig. 6I). Additionally, the perturbed cells exhibited increased rhythm irregularities, with a higher percentage of irregular beats compared to control-treated cells (Fig. 6J). Overall, overexpression of predicted novel pro-aging targets disrupted gene programs involved in limiting inflammation, preserving mitochondrial function, and restraining senescence-associated secretory factors, leading to functional defects in calcium handling and rhythm that may contribute to age-related cardiovascular decline.

### Top predicted pro-aging drivers induced cardiac dysfunction *in vivo*

Among the targets validated to drive aging-related gene network dysregulation and functional decline in cardiomyocytes, P4HA1 and RASGEF1B were predicted to have the broadest effects across cardiac cell types (Fig. 6E). Indeed, when we tested overexpression of these targets in human primary cardiac fibroblasts, we found that both perturbations again led to substantial downstream network shifts that significantly overlapped with gene network dysregulation in cardiomyocytes (Fig. 6K, S2G-H, Table S4). The downstream dysregulation shared between both targets in both cell type contexts was significantly enriched for multiple pathways relevant to age-related decline including response to stress and DNA damage/telomere stress-induced senescence (Table S6). Furthermore, both pro-aging perturbations led to significantly increased senescence in human primary cardiac fibroblasts, a hallmark of age-related dysfunction (Fig. 6L).

Finally, we tested whether overexpression of either P4HA1 or RASGEF1B in cardiomyocytes was sufficient to cause cardiovascular decline *in vivo*. To this end, we delivered AAV9 constructs encoding each factor to adult mice and assessed cardiac function six weeks post-injection (Fig. 6M). Remarkably, both perturbations led to a significant decline in cardiovascular function within six weeks of their overexpression (Fig. 6N). Global longitudinal strain and ejection fraction were both affected in echocardiographic measurements, indicating reduced systolic function in response to the perturbations.

In sum, MaxToki predicted pro-aging drivers that had a true impact on cardiovascular function in vivo, pointing to potential targets for intervention to prevent age-related cardiovascular decline.

## Discussion

Despite major advances in modeling cell states from single-cell transcriptomic data, existing computational frameworks largely treat cell states as static snapshots, limiting their ability to model continuous temporal progression or predict trajectory-shifting interventions. Here, we developed a temporal AI model, MaxToki, for generating cell states across dynamic trajectories and predicting perturbations to induce desired trajectory shifts. We trained both a 217M and 1B parameter model with dense attention on nearly 1 trillion gene tokens with an expanded context size of 16,384 to model multiple cells in time series, implementing advances in GPU acceleration to optimize throughput and memory usage. Our prompting strategy enabled querying the model for cell state forecasting as well as predicting the drivers of cell state progression over time with in-context learning of new trajectories unseen during training. The ability to generalize to unseen settings was made possible by the self-supervised training objective where the model learned from contextual trajectories of unlabeled cell states and from timelapses encoded by continuous numerical tokenization. The model therefore learned that timelapses occur on a numerical continuum rather than considering ages as categorical disconnected entities, enabling generalization to unseen cell types and ages.

Applying the model to predict pro-aging vs. rejuvenating perturbations in cardiac cell types revealed novel targets influencing aging-related gene network dysregulation and functional decline that were experimentally validated in cardiomyocytes and in vivo, demonstrating the utility of the model in driving verifiable biological insights. The predicted targets were indeed found to modulate gene programs involved in inflammation, mitochondrial dysfunction, and senescence-associated secretory factors, which are critical components of age-related cardiovascular decline. Furthermore, one of the targets, P4HA1, was previously reported to be downregulated by GLP1 agonists (*34*), opening the opportunity to use temporal AI models to predict the mechanisms of cardiovascular resilience offered by this new class of therapeutics.

Interestingly, the model also inferred age acceleration in age-related diseases not observed during training, including pulmonary fibrosis and Alzheimer dementia. Age-related diseases may involve both acceleration of normal age-dependent decline along with dysregulation along an interacting disease axis. Future work is needed to understand the interaction between these axes and whether interventions that slow age-related dysfunction will promote resilience to age-dependent diseases. Of course, one limitation in understanding the progression of diseases over long timelapses is that many diseases affect clinically inaccessible tissues where repeat sampling in the same individual over the disease course is not possible.

Accordingly, future data generation initiatives leveraging iPSC disease models may complement primary tissue data by combining time-resolved profiling with experimental perturbations to derive temporal, causal maps of cellular responses over time. Such efforts would provide instrumental data for temporal AI models to learn the causal mechanisms of progression along the disease axis and how interventions that slow normal age-related decline may protect against these age-dependent conditions.

Overall, MaxToki represents a temporal AI model for generating cell states across dynamic trajectories and predicting perturbations to induce desired trajectory shifts that can now be applied to accelerate the discovery of candidate targets for programming therapeutic cellular trajectories.

## Acknowledgements

We thank Jack Rae and the Theodoris Lab members for helpful scientific discussions. CVT was supported by grants from the Longevity Impetus Grant, Searle Scholars Program, National Institutes of Health (DP5OD036170), Burroughs Wellcome Fund Career Award for Medical Scientists (1022136.01), and Dana and Robert Hamwee. MA was supported by the National Institutes of Health grants R01HL181372 and R01HL175312, the American Heart Association Second Century Award (24SCEFIA1252167), and the Longevity Impetus Grant; and he is the Dario and Irina Sattui Investigator at Gladstone Institutes. JUW and SD were supported by the German Research Foundation (SFB1366, Project 394046768 and SFB1531, Project 456687919) and the European Research Council Advanced Grant (Neuroheart).

## Author Contributions

JGO and CVT conceived of the work. JGO developed the models and designed/performed computational analyses. RDN designed/performed in vivo experimental validation studies. RDN, JUGW, SDK, and BK designed/performed in vitro experimental validation studies. AK and KW provided data from OSKM partial reprogramming and contributed to analysis design. MMR provided data from Alzheimer cohort 2 and contributed to analysis design. SK developed code for GPU acceleration. TL contributed to model and analysis design. LZ provided the telomere-shortened mouse data. HC and YZ contributed to assembly of the pretraining corpuses. SA and WAJ contributed to model evaluation. SL contributed to interpretability analyses. CT contributed to Alzheimer cohort 2 analysis. DMP contributed to calcium cycle kinetics recordings and analysis. AJC and ACY designed analyses and supervised data generation for Alzheimer cohort 2. NT supervised code development for GPU acceleration. SY designed analyses and supervised OSKM partial reprogramming experimental studies. SD and MA designed analyses and supervised experimental validation studies. CVT designed analyses and supervised model development and experimental validation studies. JGO and CVT wrote the manuscript. All authors edited the manuscript.

## Data and Code Availability

Genecorpus-175M will be available on Hugging Face Dataset Hub. RNA sequencing data will be deposited on Gene Expression Omnibus. The pretrained MaxToki models are available on Hugging Face Model Hub (https://huggingface.co/theodoris-lab/MaxToki). The generative pretraining, timelapse training, and inference code are available at the NVIDIA Digital Biology Research GitHub repository (https://github.com/NVIDIA-Digital-Bio/MaxToki).

## Supplementary Materials

### Methods

#### Training and inference dataset assembly

##### First stage training corpus: Genecorpus-175M

We assembled a large-scale pretraining corpus, Genecorpus-175M, comprising ∼175 million human single-cell transcriptomes (173,013,164 post-filtering as described below) from a broad range of tissues from 10,795 publicly available datasets (Table S1). Cells with high mutational burdens like malignant cells and immortalized cell lines were excluded as they are known to contain gain of function variants that alter gene functions from what the model would observe in other cells with low mutational burdens. DOIs were cross-referenced between studies to ensure datasets were unique to avoid inclusion of duplicated cells within the corpus. Publicly available datasets containing raw counts were collected from National Center for Biotechnology Information (NCBI) Gene Expression Omnibus (GEO), NCBI Sequence Read Archive (SRA), CELLxGENE, scBaseCount, Human Cell Atlas, European Molecular Biology Laboratory-European Bioinformatics Institute (EMBL-EBI) Single Cell Expression Atlas, Broad Institute Single Cell Portal, Brotman Baty Institute (BBI)-Allen Single Cell Atlases, Tumor Immune Single-cell Hub (TISCH) (excluding malignant cells), Panglao Database, 10x Genomics, University of California, Santa Cruz Cell Browser, European Genome-phenome Archive, Synapse, Riken, Zenodo, National Institutes of Health (NIH) Figshare Archive, NCBI dbGap, Refine.bio, China National GeneBank Sequence Archive, Mendeley Data, and individual communication with authors of the original studies (Table S1). Additional resources for collecting information about suitable studies included Entrez Direct tools and the dataset summary from Svensson et al., Database 2020 (*35*). Tools utilized in conversion of data to uniform files included loompy, scanpy, anndata, scipy, numpy, pandas, Cell Ranger, and LoomExperiment. Gene annotation data was retrieved from Ensembl, NCBI, and HGNC (2023-11-01) databases and additionally queried through MyGene (*36*). Raw and unfiltered data files were processed to remove empty droplets and debris using STAR version 2.7.8a with the Cell Ranger2.2 (run mode – soloCellFiltered).

##### Second stage training corpus: Genecorpus-Aging-22M

Genecorpus-Aging-22M was assembled from ∼22 million human single-cell transcriptomes from the CELLxGENE corpus annotated as “normal” in the disease attribute. The assembled data represented 595 cell types across 3805 total donors aged newborn to the tenth decade of life, balanced by sex (49% male, 51% female). The “is_primary_data” flag was used to prevent collecting duplicated cells. 18 cell types were held-out from training to facilitate evaluation on unseen trajectories. These cell types were randomly chosen from those with the lowest representation of ages. Additionally, ages 21, 31, 41, 51, 61, 71, and 81 were held-out from training to facilitate evaluation on held-out ages and held-out donors.

##### Pseudotime trajectory training data

Short timelapse training data was composed of single-cell transcriptomes from partial reprogramming of primary adult human cytotoxic T lymphocytes, dermal fibroblasts, and aortic endothelial cells, obtained from 4 donors per cell type. The partial reprogramming with OCT4, SOX2, KLF4, and c-MYC (OSKM) via Sendai viral vectors represents a rejuvenating perturbation that partially reprograms cells without fully inducing pluripotency. Reprogramming was induced for 5 days for cytotoxic T lymphocytes and 7 days for dermal fibroblasts and aortic endothelial cells. At the terminal time point (day 5 for cytotoxic T lymphocytes; day 7 for dermal fibroblasts and aortic endothelial cells), cells were sorted by FACS into stage-defined populations using a somatic identity marker and a pluripotency marker. Three stages of reprogramming were defined as stage 1: positive for somatic identity marker and negative for pluripotency marker, stage 2: negative for both somatic identity marker and pluripotency marker, and stage 3: negative for somatic identity marker and positive for pluripotency marker. Somatic identity and pluripotency markers used for sorting the three cell types of cytotoxic T lymphocytes, aortic endothelial cells, and dermal fibroblasts were as follows, respectively: CD8a, SSEA4; CD13, SSEA5; and CD31, SSEA4. For aortic endothelial cells, reprogramming efficiency was low, and stage 3 cells could not be consistently obtained.

Single-cell multiome libraries were generated using the 10x Genomics Chromium Next GEM Single Cell Multiome ATAC + Gene Expression platform following the manufacturer’s protocols and sequenced on an Illumina NovaSeq X. Raw sequencing data were processed with Cell Ranger ARC v2.1.0 (10x Genomics). For the analyses in this manuscript (Fig. 3), we used only the gene expression (RNA) modality from the single-cell multiome data.

Pseudotime trajectory inference was performed using MIRA (*37*) to derive the pseudotime “ages” for each transcriptome. Default arguments were used with 60 trials, optimizing over a minimum of 3 and maximum of 20 topics. Inflection intervals between the marker-defined stage 1, 2, and 3 cells were selected to hold out from the training, as shown in Fig. 3A. Of note, the rate of change between the proportion of cells of each marker-defined stage at each interval does not directly reflect the rate of change of the expression of all genes, but only the binary detection of the indicated markers. (The full transcriptome was not utilized in defining the held-out intervals.)

##### Age acceleration inference data

Data for age acceleration inference in mucosal epithelial cells in the lung of donors exposed to heavy smoking vs. non-smoking age-matched controls was drawn from Goldfarbmuren et al, *Nature Communications* 2020 (*18*). Single-cell transcriptomes from non-smoking donors were used to construct context trajectories in the prompt. Query cells were drawn from either donors exposed to heavy smoking (T85, T120, T154, T167) or non-smoking age-matched controls (T164, T165, T166). This dataset was not part of Genecorpus-Aging-22M.

Data for age acceleration inference in lung fibroblasts from patients with pulmonary fibrosis vs. age-matched controls were drawn from Sikkema et al, *Nature Medicine* 2023 (*20*). Cells labeled as fibroblast or a subtype thereof were included except those labeled as derived from the bronchus (“bronchus fibroblast of the lung”). Single-cell transcriptomes from control donors were used to construct context trajectories in the prompt. Query cells were drawn from either donors affected by pulmonary fibrosis (three total donors available aged 40, 51 and 57) or age-matched control donors (aged 35, 52, 56, 57). Comparisons were performed with matched age groups ({35, 40}, {51, 52}, and {56, 57}). Pulmonary fibrosis samples were not part of Genecorpus-Aging-22M.

Data for age acceleration inference in microglia from Alzheimer patients vs. age-matched controls in Alzheimer dataset 1 were drawn from Mount Sinai NIH Neurobiobank, Lee et al, *Genetic and Genomic Medicine*, 2024 (*24*). Alzheimer dataset 1 analysis used cells labeled as microglia (finest annotation available from original authors in CELLxGENE). Single-cell transcriptomes from control donors were used to construct context trajectories in the prompt. Query cells were drawn from either control donors of ages excluded from MaxToki second stage training (61, 71, and 81) or age-matched donors affected by Alzheimer dementia (label: “dementia || Alzheimer disease”) from ages within two years of the control groups (59, 62, 69, 71, 79, and 81). Comparisons were performed with matched age groups ({61, 59, 62}, {71, 69}, and {81}). Specific donor IDs used were as follows: control: Donor_1052, Donor_1129, Donor_1135, Donor_1298, Donor_1301, Donor_1434, Donor_1448, Donor_1466, Donor_17, Donor_174, Donor_575, Donor_71, Donor_725, Donor_76, Donor_78, Donor_840, Donor_914, Donor_923, Donor_948, Donor_984; Alzheimer dementia: Donor_1035, Donor_1309, Donor_1433, Donor_190, Donor_30, Donor_322, Donor_482, Donor_58, Donor_65, Donor_777, Donor_909, Donor_925, Donor_953. Control and disease query cells used for this analysis were held-out from MaxToki second stage training and unseen by the model.

Alzheimer dataset 2 was composed of single-cell transcriptomes from samples from Duke and Johns Hopkins Alzheimer Disease Research Centers and the Combes and Yang Labs. Post-mortem fresh frozen dlPFC (BA9/46) samples were obtained from the Duke and Johns Hopkins Alzheimer Disease Research Centers, spanning diagnoses from no cognitive impairment to mild cognitive impairment, Alzheimer disease, and cognitive resilience. Vascular and parenchymal nuclei were isolated and pooled using a dextran-based density centrifugation protocol followed by enzymatic digestion, as previously described (*38*). Nuclei were fixed, sorted by flow cytometry, and processed using the 10x Genomics Chromium Fixed RNA Profiling Kit (Flex, PN-1000414). Libraries were sequenced on the Illumina NovaSeqX at a target depth of ∼40,000 reads/cell. Raw reads were aligned to the human GRCh38 reference genome and demultiplexed using Cell Ranger v7.0.1. Ambient RNA and doublets were removed using CellBender v0.3.0 and DoubletFinder v2.0.3. Low-quality cells were filtered on minimum unique gene count and percent mitochondrial reads, and libraries were integrated using Seurat v4.3.0 with Harmony batch correction.

Because this dataset was composed of 10X Flex data that is a probe-based technology causing systematic biases in the quantified RNA abundance compared to the predominant 10X technologies in the pretraining dataset, MaxToki was first fine-tuned on control samples from this dataset, holding out a subset of control samples and all Alzheimer, mild cognitive impairment, and resilient (same neuropathology, no cognitive impairment) samples for the age acceleration inference analysis. The aging-trained MaxToki was fine-tuned with prompts that included context trajectories from 59 total control donors ranging from ages 21 to 100 and query cells from a subset of these control donors (15 total) aged 64 to 100. Fine-tuning was performed with 1 epoch on 6400 example trajectories with bf16 mixed precision, effective batch size of 256, AdamW optimizer, cosine learning schedule, maximum learning rate of 3.19e-5, weight decay of 6.32e-2, and warmup ratio of 7.81e-3. Validation was performed in held-out control donors aged 67, 74, and 84 to confirm model generalizability. Aging acceleration inference studies were performed using this fine-tuned model comparing the predicted timelapses for homeostatic microglia in unseen age-matched controls (aged 66, 87, 87), donors affected by Alzheimer (aged 64, 72, 76, 77, 79, 81, 83, 88, 90, 90, 91, 92), donors with mild cognitive impairment (aged 76, 78, 81, 84, 86, 93), resilient donors (aged 73, 74, 79, 80, 82, 84, 85, 86, 91). This dataset was not part of Genecorpus-Aging-22M.

#### Rank value encoding of single-cell transcriptomes

Each transcriptome was presented to the model as a rank value encoding as previously described (*3*). The rank value encodings are a nonparametric representation of the transcriptome that takes advantage of the many observations of the gene’s expression across the entire Genecorpus to prioritize genes that distinguish cell state. Specifically, this method will deprioritize ubiquitously highly-expressed housekeeping genes by normalizing them to a lower rank. Conversely, genes such as transcription factors that may be lowly expressed when they are expressed but highly distinguish cell state will move to a higher rank within the encoding.

Furthermore, this rank-based approach may be more robust against technical artifacts that may systematically bias the absolute transcript counts value while the overall relative ranking of genes within each cell remains more stable.

The rank value encodings were constructed as previously described (*3*). The scaling factor for each gene was derived from the non-zero median value of expression of each detected gene across all cells in the pretraining corpus passing quality filtering that were sequenced on droplet-based platforms, excluding cells with high mutational burdens such as malignant cells and immortalized cell lines. After scaling the expression of each gene, the genes were ordered by the rank of their scaled expression in that specific cell. The rank value encoding for each single-cell transcriptome was then tokenized on the basis of a vocabulary of 20,271 protein coding genes detected within the pretraining corpus. The vocabulary also included four special tokens: a padding, masking, BOS (beginning of state), and EOS (end of state) token, for a total vocabulary size of 20,275. A BOS and EOS token were added to the beginning and end of each rank value encoding, respectively. The tokenized dataset was stored within the Hugging Face Datasets structure, which is based on the Apache Arrow format that allows processing of large datasets with zero-copy reads without memory constraints.

Of note, this strategy is also space-efficient as the genes are stored as ranked tokens as opposed to the exact transcript values, and we only store genes detected within each cell rather than the full sparse dataset that includes all of the undetected genes. This also prevents wasting computation on zeros, as the model learns from the absence of genes from a rank value encoding without having to explicitly instruct the model that they have zero expression. This is analogous, for example, to how natural language models learn that a statement may have “positive” meaning based on the absence of “negative” words, without needing to present the remainder of the absent words from the natural language dictionary at the end of every sentence to explicitly instruct the model they are not present.

#### Temporal encoding of cell state trajectories

Temporal encodings were composed of multiple cells in series, each bounded by the BOS and EOS token as in the pretraining corpus, with the addition of a numerical timelapse token between each cell indicating the time elapsed between the given states along the trajectory. The prompt structure involved an initial context trajectory (generally 2-3 cell states and the intervening timelapses between them) followed by a query bounded by special query tokens

BOQ (beginning of query) and EOQ (end of query). The content of the query was either (task 1) a query cell, prompting the model to respond with the numerical time token indicating the time elapsed between the last context cell and the query cell, or (task 2) a timelapse, prompting the model to respond with generating the rank value encoding for the cell transcriptome that would occur after the given time was elapsed from the last context cell. For example, a prompt with a 2-cell context trajectory would be constructed as follows:

Prompt for task 1: [context cell 1] [timelapse 1] [context cell 2] [BOQ] [query cell] [EOQ] Expected response for task 1: [response timelapse]

Prompt for task 2: [context cell 1] [timelapse 1] [context cell 2] [BOQ] [query timelapse] [EOQ] Expected response for task 2: [response cell]

Trajectories were constructed with cell states from the same cell type and sex for the aging applications and with cells from the same cell type and donor for the partial reprogramming pseudotime application. The cell type, sex, and donor information were not explicitly provided to the model; instead, the model was compelled to learn from the context trajectory in the prompt to generate responses specific to the given context.

#### MaxToki architecture and two-stage training strategy

##### MaxToki architecture

MaxToki is composed of dense transformer decoder units, each composed of a self-attention layer and feed forward neural network. The input size for the first stage training was 4096 genes per cell, which fully represents 93% of the cells in Genecorpus-175M when considering genes within the model vocabulary of 20,271 protein coding genes. RoPE was used for position embeddings to enable extending the context length in the second stage training, which models multiple cells along temporal trajectories.

Two model parameter sizes were trained, a 217 million parameter model and a 1 billion parameter model. The 217 million model parameters were as follows: 11 layers, 1232 embedding dimensions, 8 attention heads, and 2464 feed forward size. The 1 billion model parameters were as follows: 20 layers, 2304 embedding dimensions, 16 attention heads, and 4608 feed forward size. Modeling was implemented in pytorch and using Hugging Face Transformers and NVIDIA BioNeMo, NeMo, Megatron-LM, Megatron-Core, and Transformer Engine for model configuration, data loading, and training.

##### MaxToki first stage training: cell state generation

MaxToki was first trained with an autoregressive objective to learn to generate single-cell transcriptomes based on the rank value encodings from ∼175 million (173,013,164) single-cell transcriptomes from Genecorpus-175M (excluding cells with high mutational burdens such as malignant cells and immortalized cell lines). The model was tasked with generating the next gene within the rank value encoding given the context of the preceding genes. A major strength of this approach is that it is entirely self-supervised and can be accomplished on completely unlabeled data, which allows the inclusion of large amounts of training data without being restricted to samples with accompanying labels. Pretraining hyperparameters were optimized to the following final values: max learning rate: 0.00016; learning scheduler: cosine with warmup; optimizer: Adam with weight decay fix; warmup ratio: 0.00735; effective batch size (batch size x GPUs): 256 for 1B model, 144 for 217M model. Weights & Biases was used for experimentation tracking.

As the input size of 4096 is considerably large for a full dense self-attention model and transformers have a quadratic memory and time complexity 𝒪*(L^2^)* with respect to input size, we implemented the following strategies for GPU acceleration. The 217 million parameter model was trained in full precision (fp32) using distributed GPU training algorithms (*39*, *40*) to allow efficient pretraining on large-scale data using Deepspeed. This approach partitions parameters, gradients, and optimizer states across available GPUs, offloads processing/memory as possible to CPU to allow more to fit on GPU, and reduces memory fragmentation by ensuring long and short term memory allocations do not mix. Training was distributed across 24 Nvidia H100 80GB GPUs.

The 1 billion parameter model was trained with mixed precision (bf16) using FlashAttention-2 and Transformer Engine, optimized using components of the NVIDIA BioNeMo stack, built on NeMo, Megatron-LM, Megatron-Core, and Transformer Engine. To enable FlashAttention-2, we modified model parameter shapes so that feed-forward hidden dimensions were compatible with attention head partitioning by ensuring the feed-forward hidden dimensions evenly divided the number of attention heads. We then activated FlashAttention-2 through the integrated stack, leveraging fused kernels across NeMo, Megatron-LM, and Transformer Engine. Training was distributed across 32 Nvidia H200 141GB GPUs.

##### MaxToki second stage training: temporal training and prompting strategy

In the second stage training, MaxToki learned to generate cell states and the timelapses between them across dynamic trajectories. The context length was extended to 16,384 with RoPE scaling to accommodate multiple transcriptomes within the input prompt. RoPE scaling modifies the Rotary Positional Embeddings of the model by reducing the frequency of rotation to interpolate more tokens into the existing positional framework (*16*). The token dictionary was also expanded to include the timelapse tokens and special tokens for the beginning and end of query (BOQ and EOQ, respectively). The second stage training was performed with mixed precision (bf16) and an effective batch size of 144 and 128 for the 217M and 1B model, respectively; otherwise hyperparameters were consistent with the first stage training.

MaxToki was prompted with a temporal encoding as discussed above in *Temporal encoding of cell state trajectories*, which includes an initial context trajectory (generally 2-3 cell states and the intervening timelapses between them) followed by a query. The query consisted of either (task 1) a query cell, prompting the model to respond with the numerical time token indicating the time elapsed between the last context cell and the query cell, or (task 2) a timelapse, prompting the model to respond with generating the rank value encoding for the cell transcriptome that would occur after the given time was elapsed from the last context cell. The loss was calculated only for the model response by dynamically detecting the EOQ (end of query) token and masking the loss of the prompt up to and including EOQ.

Mean squared error (MSE) loss was used for task 1 to promote the model to learn that the timelapses fall along a numerical continuum, while cross-entropy (CE) loss was used for task 2. Task 1 either used the final hidden state to predict the timelapse with a separate MSE regression head (for the 217M model) or directly quantified MSE based on the output logits for the generated response (for the 1B model). For the latter method, we mapped the output logits to a continuous scalar by calculating the expected value across the numerical token distribution. Specifically, we computed the weighted average of the token values based on their softmax-normalized probabilities. The resulting value was then compared to the ground-truth value using MSE. Additionally, a penalty was applied when the model generated a non-timelapse token in the response to quickly encourage the model to respond with a timelapse to task 1 queries. The penalty factory was a tunable parameter for which we used 100.

For inference, we adopted the Megatron-Core inference engine using the DynamicInferenceContext abstraction. This object manages key–value cache allocation and reuse across decoding steps and supports sequence packing for efficient utilization of compute and memory. These optimizations resulted in a >400x improvement in autoregressive generation speed relative to the baseline implementation. We also implemented restrictions for the model to generate a maximum of 4096 tokens per cell response and no repeated genes within a single cell.

The evaluation, interpretability analysis, age acceleration, and in silico perturbation experiments reflect the smaller 217 million parameter model to optimize efficiency under resource constraints and demonstrate capabilities of the more accessible model size.

#### Evaluation on held-out ages and cell types

##### Evaluation of timelapse predictions

Timelapse predictions were evaluated on ages held-out from the training (21, 31, 41, 51, 61, 71, 82), which also represented held-out donors. The query cell provided within the trajectory was of a held-out age and the model was tasked with producing the timelapse between the last context cell and the query cell of the held-out age. The error of the MaxToki-predicted timelapse vs. the ground truth timelapse was quantified and compared to a linear model (SGDRegressor, see below in *Comparison to linear model*) or the baseline error of selecting the timelapse that would correspond to the query cell being the most common age for the given cell type and sex in the training data. The correlation of the predicted timelapse to the ground truth timelapse was also quantified, including for trajectories separated into those where the query cell was a past, intervening, or future cell state with respect to the other cell states within the trajectory to quantify relative performance in these three domains.

An analogous analysis was performed with cell types held-out from the training (enteroendocrine cell of colon, myeloid dendritic cell, tuft cell of colon, mesothelial fibroblast, epithelial cell of small intestine, epithelial cell of thymus, keratocyte, tongue muscle cell, colon goblet cell, mature microglial cell, ventricular cardiac muscle cell, eye photoreceptor cell, cortical thymic epithelial cell, early T lineage precursor, IgG-negative class switched memory B cell, mature astrocyte, IgG memory B cell, radial glial cell). The full trajectory in the prompt was composed of a given held-out cell type, requiring in-context learning for the model to understand time progression within that cell type’s aging trajectory to predict the timelapse between the query cell and the last context cell. Again, the error of MaxToki-predicted timelapse vs. the ground truth timelapse was quantified and compared to the baseline error of selecting the timelapse that would correspond to the query cell being the most common age for the given cell type and sex, in the test data itself. The correlation of the predicted timelapse to the ground truth timelapse was also quantified.

##### Comparison to linear model

MaxToki was also compared to a linear model, the sklearn implementation of SGDRegressor. SGDRegressor was chosen due to memory constraints because it processes data incrementally, rather than needing to load the entire dataset into RAM. For this comparison, SGDRegressor was compared to MaxToki while training both on ∼4M examples. For SGDRegressor, the same examples were converted to a compatible format by extending each cell transcriptome in the prompt to a vector of all genes in the token dictionary where the value for each gene was (4096 minus the rank)/4096, where the rank refers to the gene’s rank within the rank value encoding provided to MaxToki. All undetected genes were assigned a rank of 0. The scaling to 0 to 1 was necessary to stabilize the model training. The values for the timelapses were included in the vector in the same positions as for the MaxToki examples, directly providing their numerical value, scaled to be within -1 and 1, again for model stabilization. The SGDRegressor was then trained using a batch size of 5000 and otherwise default settings. For comparisons with SGDRegressor, both SGDRegressor and MaxToki were trained with ∼4M example trajectories.

##### Evaluation of generated cells

The generated cells for held-out ages were evaluated based on concordance with the age of their nearest ground truth neighbor within the embedding space of an external model, Geneformer (V2-316M variant) (*3*). Geneformer was first fine-tuned for a single epoch on 480,000 single-cell transcriptomes from Genecorpus-Aging-22M, excluding the same held-out ages/donors as the MaxToki training, to classify cells based on their age bin: infant (0-1 year), toddler (1-5 years), child (5-10 years), teen (10-20 years), 20’s, 30’s, 40’s, 50’s, 60’s, 70’s, 80’s, and 90+, with a max of 40,000 training cells per bin. Hyperparameters were optimized over 25 trials with 10 frozen layers, batch size of 56, cosine learning rate scheduler, learning rate 1e-5 to 1e-4, weight decay 0.01 to 0.3, and warmup ratio 0.005 to 0.05 (remainder default). The ground truth vs. generated transcriptomes for the held-out ages were then evaluated within the Geneformer cell embedding space. The age of the nearest ground truth neighbor for each generated transcriptome was identified, and correlation between the prompted age (based on the query timelapse and last context cell) and nearest ground truth age was quantified.

The generated cells were also evaluated for their concordance with the cell type in the prompted trajectory. Firstly, the annotations of generated vs. ground truth cells by an external cell type classifier, based on the Geneformer model (V2-104M variant) (*3*), were compared.

Geneformer was first fine-tuned to classify cell types based on uniform annotations within the CELLxGENE corpus with the following optimal hyperparameters selected from 15 trials: max learning rate: 2e-05; learning scheduler: cosine with warmup; optimizer: Adam with weight decay fix; warmup ratio: 0.09; weight decay: 0.01; batch size: 64; max gradient normalization: 1.3; dropout rate: 0.3; frozen layers: 7; epoch: 1. The MaxToki generated vs. ground truth cells for held-out ages (which were also excluded from the Geneformer fine-tuning) were then annotated by the Geneformer cell type classifier and concordance of their annotations was quantified. The embeddings of the MaxToki generated and ground truth cells held-out from the Geneformer training were also visualized by UMAP alongside ground truth cells seen by the Geneformer model to test for co-embedding of each cell type.

Secondly, the MaxToki generated cells were evaluated for their ability to newly train a cell type classifier to annotate unseen ground truth cells. The pretrained Geneformer model (V2-316M variant) (*3*) was fine-tuned with either 1) ground truth cells, 2) ground truth cells with shuffled genes within the rank value encoding (“bag of genes” cells), or 3) MaxToki generated cells for held-out ages and tested for its ability to correctly annotate held-out ground truth cells. Only if the generated cells retained key biological informational content for cell type identity would an external model trained only on the generated cells be able to classify the cell type of unseen ground truth cells. All models were fine-tuned with a max of 4000 cells per class (epithelial cell, myeloid cell, T cell, neuron, B cell, fibroblast, vascular endothelial cell, endocrine cell, hematopoietic progenitor, interneuron, hematopoietic progenitor, mural cell, ciliated cell, erythroid, smooth muscle cell). Hyperparameters were optimized over 10 trials with 10 frozen layers, batch size of 56, cosine learning rate scheduler, learning rate 1e-5 to 1e-4, weight decay 0.01 to 0.3, and warmup ratio 0.005 to 0.05 (remainder default).

Finally, the MaxToki generated cells were evaluated for their resolution. We first trained Geneformer (V2-104M variant) (*3*) to distinguish single cells from simulated doublets derived by merging two cells of the same fine cell type from cells in the CELLxGENE corpus. Specifically, doublets were simulated by randomly selecting two cells of the same fine cell type, interlacing their rank value encodings, deduplicating the genes via random selection of one gene copy to keep, and then truncating to the average number of genes between the two cells. Fine-tuning was performed with 8 frozen layers, batch size 8, gradient accumulation steps 4, learning rate for pretrained layers 5e-5, learning rate for classifier head 1e-3, warmup ratio 0.1, weight decay 0.01, dropout rate 0.1, maximum gradient norm for clipping 1.0, seed 37. The trained doublet detector model was then applied to classify the resolution of held-out ground truth cells and simulated doublets of the same or different cell types as well as the MaxToki generated cells.

#### Pseudotime training and evaluation

##### Pseudotime trajectory inference and MaxToki training

The pretrained 217M MaxToki model that had undergone the first stage training was then subjected to a second stage temporal training using the partial reprogramming data, constructed as temporal encodings with timelapses in units of MIRA-inferred pseudotime as discussed above. The token dictionary for this training was thus expanded to include the timelapse tokens relevant to the pseudotime units (along with special tokens for the beginning and end of query, BOQ and EOQ, respectively). The training was otherwise performed as discussed above in *MaxToki second stage training: temporal training and prompting strategy* for a total number of 1,697,280 example trajectories, reflecting the optimal CE loss for cell generation.

##### Evaluation on held-out inflection intervals

The model was then evaluated on its performance in both task 1 and 2 for query cells or timelapses reflecting the held-out inflection intervals. Pseudo-timelapse prediction for query cells in held-out intervals (task 1) was compared to the baseline of predicting the pseudo-timelapse to the most prevalent pseudotime for the given cell type and donor in the prompted trajectory. The ground truth pseudotime of the query cells was compared to the inferred pseudotime of the query cells, derived by adding the predicted pseudo-timelapse to the ground truth pseudotime of the last context cell.

The ground truth cells within held-out inflection intervals were then compared to the cells generated in response to a query pseudo-timelapse that would reflect a ground truth pseudotime position within the held-out inflection intervals. Cytotoxic T lymphocytes were chosen for this analysis due to having the greatest pseudotime separation between the held-out inflection intervals. Genes were tested for upregulation or downregulation by rank using the position within the generated or ground truth rank value encoding and determining the differential ranking between the control and experimental conditions (i.e. the change in the gene’s rank between the generated or ground truth cells within held-out inflection interval 2-3 compared to the other cells within the trajectory). Specifically, the ranks were processed as follows: the rank normalized by the total sequence length of that cell was subtracted from 1 and then multiplied by 4095 and added to 1 so that every detected gene in the cell had a minimum value of 1 and maximum value of 4096 (the maximum sequence length) and undetected genes were assigned a value of 0. These normalized rank values were then processed as pseudo-counts with standard Scanpy methods: normalized to a total of 10,000, log1p transformed, and compared using rank_gene_groups to obtain the final log2 fold change (differential ranking) between conditions. Correlation was quantified between the differential ranking for each gene in the generated cells and the differential ranking for each gene in the ground truth cells.

The ground truth differential expression between the cells in held-out inflection interval 2-3 compared to other cells in the trajectory was also quantified by transcript counts using Scanpy’s standard recommended procedure and default arguments for differential expression analysis. The ground truth differential expression of the top 100 upregulated genes by rank and top 100 downregulated genes by rank either in the generated or ground truth cells was then plotted vs. the ground truth differential expression for all genes. For example, the top 100 upregulated genes by rank in the generated cells for the held-out inflection intervals 2-3 were found to be non-monotonically activated more than other genes within the cells from those intervals by ground truth differential expression of transcript counts. A similar pattern was observed for the top 100 upregulated genes by rank in the ground truth cells.

#### Age acceleration inference

Age acceleration was quantified based on whether the model inferred that more or less time had elapsed between the last context cell and a query cell that was either 1) a control cell (no known disease) or 2) an age-matched cell from a donor affected by the given condition (heavy smoking, pulmonary fibrosis, Alzheimer, mild cognitive impairment, or Alzheimer resilience). The age-matched control vs. affected prompts were constructed by matching age groups as indicated above in *Training and inference dataset assembly.* If the time elapsed was inferred to be longer than the true age difference of the samples, then the query cell was interpreted by the model to have accelerated aging.

#### Interpretability analysis

##### Ablation studies

Ablation studies were performed by either replacing the rank value encodings in the held-out cell type prompts with 1) “bag of genes” cells where the gene rank order was randomly shuffled while maintaining the presence of the same genes or 2) random genes. The error of the prediction of the timelapse between the last context cell and query cell was quantified.

Additionally, ablation studies were performed for held-out ages comparing the error of the prediction of the timelapse when the context trajectory and query cell were intact in the prompt vs. if either of those two components were masked (replaced with mask token).

##### Attention analysis

Attention analyses were performed by extracting attention weights for each attention head across each model layer for each position along the prompt sequence for 1000 random prompts for held-out ages. Prompts with two-cell context trajectories were used for the aggregate analyses in Fig. 4C, E, and F for simplicity (Fig. 4D showed three specific example prompts with three-cell context trajectories). Attention to transcription factors (*41*) vs. other genes was quantified across all examples for each individual attention head. The mean attention to each prompt component (context cell 1, time token, context cell 2, and query cell) was also quantified across all 1000 example prompts. The analysis for cell type context-specificity was performed with 56 cell types present with 10 or more example trajectories within the 1000 example prompts. If a given attention head paid significantly higher attention to a specific prompt component in one cell type compared to the others, that component in that attention head was labeled as cell type-specific. The cell types evaluated included: megakaryocyte_progenitor_cell, professional_antigen_presenting_cell, monocyte, epithelial_cell_of_lower_respiratory_tract, inflammatory_macrophage, retinal_ganglion_cell, basophilic_erythroblast, lymphocyte, kidney_distal_convoluted_tubule_epithelial_cell, memory_B_cell, pulmonary_artery_endothelial_cell, CD16-negative,_CD56-bright_natural_killer_cell, central_memory_CD4-positive,_alpha-beta_T_cell, peripheral_blood_mononuclear_cell, lung_goblet_cell, tracheobronchial_serous_cell, effector_CD8-positive,_alpha-beta_T_cell, macrophage, effector_memory_CD4-positive,_alpha-beta_T_cell, vascular_associated_smooth_muscle_cell, OFFx_cell, keratinocyte, erythroid_lineage_cell, granulocyte_monocyte_progenitor_cell, renal_principal_cell, invaginating_midget_bipolar_cell, type_II_pneumocyte, mucosal_invariant_T_cell, epithelial_cell_of_alveolus_of_lung, chondrocyte, glycinergic_amacrine_cell, ON_parasol_ganglion_cell, Mueller_cell, hematopoietic_oligopotent_progenitor_cell, CD8-positive,_alpha-beta_regulatory_T_cell, smooth_muscle_cell, natural_killer_cell, urothelial_cell, stratified_epithelial_cell, granulocyte, renal_alpha-intercalated_cell, megakaryocyte-erythroid_progenitor_cell, CD8-positive,_alpha-beta_memory_T_cell,_CD45RO-positive, central_memory_CD8-positive,_alpha-beta_T_cell, epithelial_cell, CD1c-positive_myeloid_dendritic_cell, large_pre-B-II_cell, retinal_bipolar_neuron, tracheobronchial_smooth_muscle_cell, kidney_loop_of_Henle_epithelial_cell, kidney_loop_of_Henle_thin_descending_limb_epithelial_cell, hematopoietic_stem_cell, endothelial_cell_of_artery, melanocyte, fibroblast, CD14-positive_monocyte.

#### MaxToki in silico perturbation strategy

The in silico perturbation strategy involves perturbing the genes within context or query cells within the prompt and quantifying the impact on the model’s response, either the predicted timelapse for task 1 or the generated cell for task 2. In silico perturbation analysis was applied to cardiac cell types from CELLxGENE (atrial cardiomyocytes, ventricular cardiomyocytes, endocardial cells, capillary endothelial cells, pericytes, fibroblasts, and macrophages annotated as heart tissue) by constructing temporal trajectories for each cell type and performing in silico inhibition of each gene within the query cell by removing the gene from the rank value encoding. The response of the model for task 1 (the predicted timelapse between the query cell and the last context cell) was quantified for both the unperturbed query cell and its perturbed analog while keeping the remainder of the prompt context trajectory unchanged. The difference in the predicted timelapses was quantified for each controlled pair of perturbed vs. unperturbed trajectories. Perturbations that led to the model inferring less time had elapsed (negative difference in time) indicated that the model interpreted this perturbation as rejuvenating, whereas perturbations that led to the model inferring that more time had elapsed (positive difference in time) indicated that the model interpreted the perturbation as age-promoting. Gene set enrichment analysis of genes with statistically significant impact on the timelapse by in silico perturbation was performed with GProfiler (*42*) using the default parameters but providing a background set of all genes detected in the given cell type (e.g. all cardiac fibroblast genes for the cardiac fibroblast in silico perturbation analysis).

The predicted impact of in silico inhibition of each gene was compared to the slope of the ground truth expression change of that gene across human aging. Ground truth expression was quantified as the average transcript counts for that gene across all cells from donors whose age was within one of four age bins, 1 month to 21 years old, 22-42 years old, 43-63 years old, and 64-85 years old, normalized by the total transcript counts per cell and total transcript counts for the given age bin to control for varying read depth per cell and donor, respectively. The slope of the linear expression change of each gene across the age bins was calculated.

The predicted impact of in silico inhibition of each gene was also compared to the expression change in telomere-shortened generation 4 *Tert^-/-^*mice compared to wild-type controls based on heart single-nucleus RNA sequencing data from three 15-29 week old mice in each group. Heart collection and single-nucleus RNA sequencing was performed as previously described in Stilz et al, *Eur. Heart J.* 2026 (*31*). Briefly, hearts were collected by perfusing with DPBS in vivo, snap-freezing in liquid nitrogen, mincing with pre-filtered homogenization buffer, and sorting to separate positive nuclei from cell debris. 10X single-nucleus RNA sequencing library preparation, sequencing, and data pre-processing was performed as previously described (*31*). Differential expression was quantified by Scanpy using “rank_genes_groups” with default parameters.

#### Human primary cardiac capillary endothelial cell experimental validation

Human cardiac capillary endothelial cells were purchased from PromoCell (C-12285) and cultured in endothelial basal medium (EBM, CC-3121, Lonza) supplemented with 10% fetal bovine serum (10100147, Gibco). ZBTB16 repression was mediated using the On-Target plus SMART pool siRNAs against human ZBTB16 (L-018719-00-0005, Dharmacon / Horizon).

AllStars Negative Control siRNA (1027280, Qiagen) served as negative control. Both siRNA pools (40 nM) were transfected in OptiMEM (31985070, Gibco) and Lipofectamin RNAiMAX (13778100, Invitrogen). Senescence-associated β-galactosidase was performed using the CellEvent™ Senescence Green Detection Kit (C10850, ThermoFisher) according to the manufacturer’s instructions. The β-galactosidase positive areas were quantified using ImageJ and Volocity 7 (Quorum Technologies Inc.).

#### Cardiomyocyte experimental validation

WTC11 iPSCs were differentiated into cardiomyocytes using the STEMDiff ventricular cardiomyocyte kit (#05010) using the manufacturer-provided protocols. Differentiation efficiency was quantified by cTnT staining by flow cytometry and confirmed to be >70% for each round of differentiation. Cardiomyocytes at day 17 were treated with AAV6 overexpression of either GFP or the MaxToki-predicted pro-aging targets ATF3, EYA4, NR4A3, NXN, P4HA1, or RASGEF1B. AAV6 was selected based on testing the toxicity and infection efficiency of AAV1, 2, 6, and 9 at low (10,000) and high (50,000) multiplicity of infection (MOI) (Fig. S2B). Coding sequences of interest were cloned downstream of a constitutive CAG promoter and upstream of a Woodchuck Hepatitis Virus Posttranscriptional Regulatory Element (WPRE) to enhance transgene expression. AAV6 vectors were synthesized, packaged, and titered commercially by VectorBuilder. Cardiomyocytes were transduced at MOI 10,000 based on cell count at time of cell plating, and media was changed after 48 hours. Cells were then collected at day 21 for bulk RNA sequencing via Plasmidsaurus with four replicates of each condition. Differential expression was quantified by pydeseq2 using default parameters. Each intended AAV overexpression target was confirmed to be significantly overexpressed in the treated cells.

Principal component analysis (PCA) and hierarchical clustering of the bulk RNA sequencing was performed to compare the transcriptional effects of each perturbation. Gene set enrichment analysis with GProfiler (*42*) using the default parameters with the foreground set being genes statistically significantly changed in the perturbed cells and the background set being all cardiomyocyte genes detected in the bulk RNA sequencing assay.

iPSC-derived cardiomyocytes were functionally characterized using Fluo-4 AM dye to evaluate calcium cycling kinetics and to quantify the rate of irregular beats. Cells were incubated with 5uM Fluo-4 AM for 30 minutes at 37oC in Tyrode’s solution containing 0.02% Pluronic F-127, then washed and allowed to de-esterify. After dye loading, fluorescence imaging was performed on spontaneously beating cells in dye-free Tyrode’s solution on an inverted Zeiss epifluorescence microscope equipped with a 488nm excitation source and a 510-550 nm emission filter. Time-series images were acquired at 20-50 frames per second with exposure times of 10-20ms, which allowed accurate resolution of calcium kinetics. Regions of interest were manually defined over individual cells. The time to peak and Decay Time Constant (Tau) for each calcium cycle were quantified for the control vs. experimental conditions. The proportion of irregular beats was also quantified.

#### Cardiac fibroblast experimental validation

Primary human cardiac fibroblasts (NHCF-V, CC-2904, Lonza) were cultured in cardiac fibroblast growth medium (FGM3, CC-4526, Lonza). Cells were cultured for three passages (1:2 split ratio) and were plated at cell count of 50,000 per well. The following day, fibroblasts were transduced at MOI 25,000 with AAV6 overexpression of either GFP or the MaxToki-predicted pro-aging targets P4HA1 or RASGEF1B. Cells were then collected after 72 hours for bulk RNA sequencing via Plasmidsaurus or for senescence-associated β-galactosidase assay with four replicates of each condition. Differential expression was quantified by pydeseq2 using default parameters. Each intended AAV overexpression target was confirmed to be significantly overexpressed in the treated cells. Gene set enrichment analysis with GProfiler (*42*) using the default parameters with the foreground set being genes statistically significantly changed in the perturbed cells and the background set being all fibroblast genes detected in the bulk RNA sequencing assay. Senescence-associated β-galactosidase was performed using the CellEvent Senescence Green Flow Cytometry Assay Kit (C10840, ThermoFisher) according to the manufacturer’s instructions. Percentage of cells positive for β-galactosidase was quantified with the Attune NxT flow cytometer (ThermoFisher).

#### In vivo validation of predicted cardiac pro-aging drivers

Overexpression of candidate genes in mice was achieved via AAV9-mediated transduction delivered by retro-orbital injection to 6–8-week-old adult male C57BL/6J mice at a dose of 1×10¹² genome copies (gc) per 25 g mouse. Candidate gene or GFP cDNA sequences were cloned downstream of a cardiac troponin T (cTnT) promoter to drive cardiomyocyte-specific expression and upstream of a woodchuck hepatitis virus posttranscriptional regulatory element (WPRE) to enhance transgene expression. Viral packaging was performed commercially (VectorBuilder). Cardiac function was assessed by transthoracic echocardiography as previously described (*43*). Briefly, mice were anesthetized with 1–2% isoflurane and imaged using a Vevo 3100 high-resolution imaging system (FUJIFILM VisualSonics) equipped with an MX550S probe. Measurements were acquired from B-mode and M-mode images in standard parasternal long- and short-axis views. Global longitudinal strain (GLS) was quantified using the VevoStrain analysis software. All image acquisition and analysis were performed in a blinded manner. Images of insufficient quality for accurate quantification were excluded in a blinded manner based on predefined criteria, including failure of automated speckle-tracking algorithms to reliably track myocardial wall segments during GLS analysis. Statistical analyses were performed using one-way ANOVA with correction for multiple comparisons. All animal protocols were approved by the Institutional Animal Care and Use Committees at the University of California San Francisco and in accordance with the National Institutes of Health Guide for the Care and Use of Laboratory Animals.

---

**Fig. S1.**
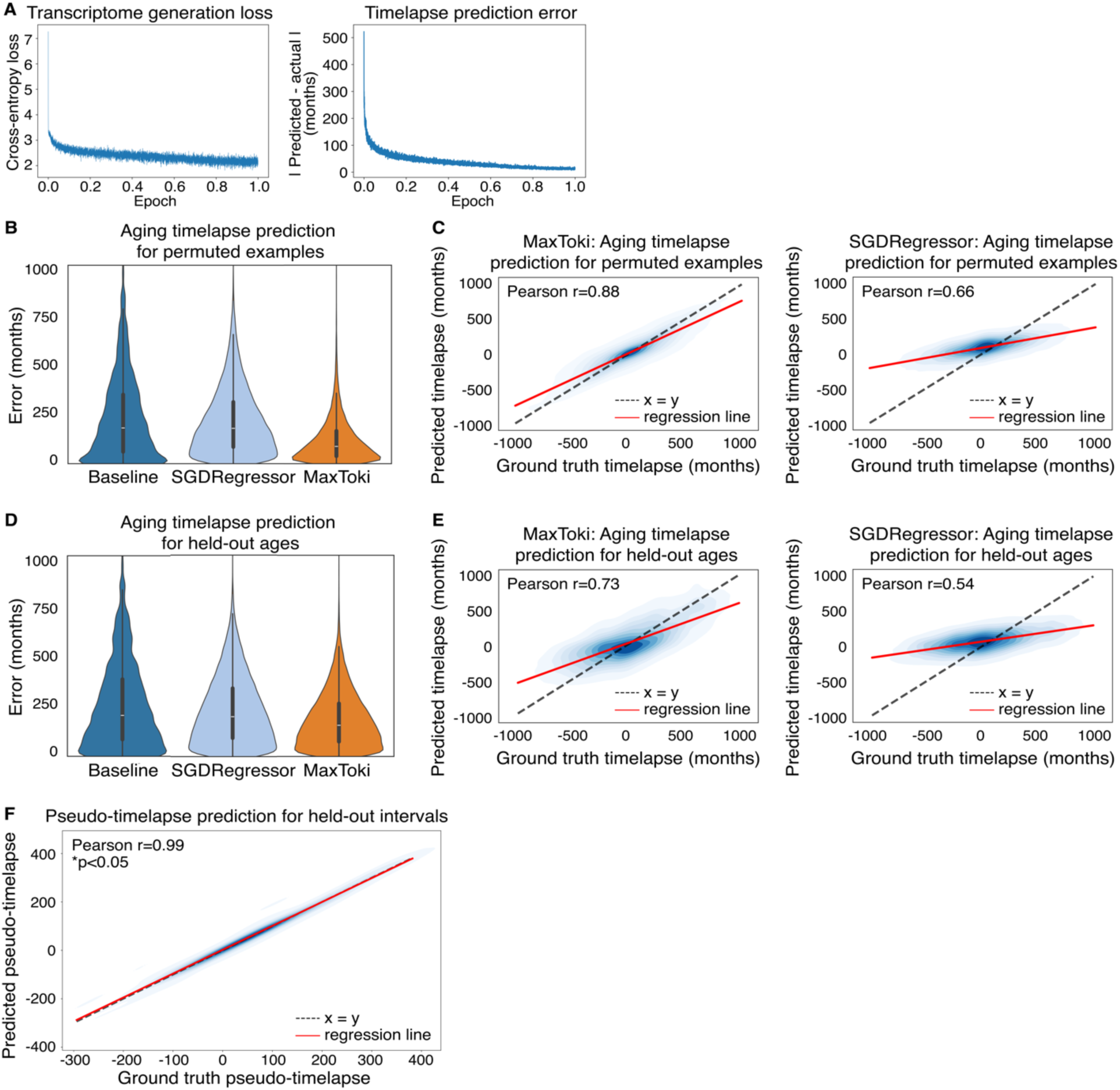
Timelapse predictions compared to alternative linear model. (**A**) Stage 2 loss curves for cell state generation (CE loss) and timelapse prediction (MSE loss) for MaxToki aging trajectory training. (**B**) Aging timelapse prediction for permuted examples by MaxToki or an alternative linear model approach (SGDRegressor) compared to baseline approach of predicting timelapses by assuming the query cell was the most common age of that cell type and sex (n=48,218). (**C**) Pearson correlation of predicted vs. ground truth aging timelapses for permuted examples for MaxToki or an alternative linear model approach (SGDRegressor) (n=48,218). (**D**) Aging timelapse prediction for held-out ages compared by MaxToki or an alternative linear model approach (SGDRegressor) to baseline approach of predicting timelapses by assuming the query cell was the most common age of that cell type and sex (n=50,113). (**E**) Pearson correlation of predicted vs. ground truth aging timelapses for held-out ages for MaxToki or an alternative linear model approach (SGDRegressor) (n=50,113). (**F**) Ground truth vs. predicted pseudo-timelapse predictions for held-out intervals in partial reprogramming application by MaxToki. In (B-E): MaxToki and SGDRegressor were trained with ∼4M example trajectories.

**Fig. S2.**
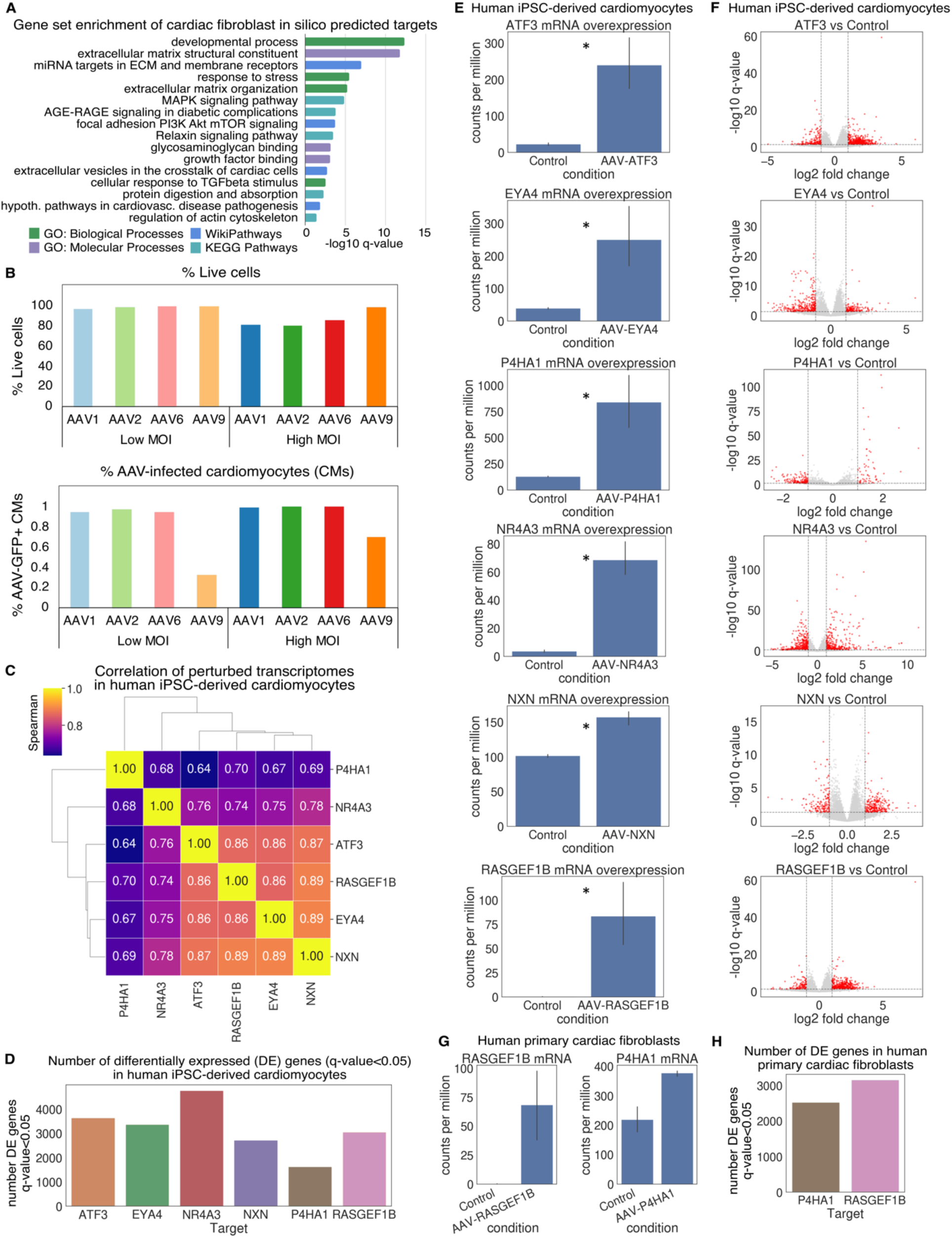
impact of predicted pro-aging perturbations in human iPSC-derived cardiomyocytes. (**A**) Gene set enrichment of in silico predicted age-modulating targets in cardiac fibroblasts (hypergeometric test with g:Set Counts and Sizes (g:SCS) correction). (**B**) % live cells and % AAV-infected iPSC-derived cardiomyocytes (percentage of the cTNT+ cells that are also AAV-GFP+) with the indicated AAVs as quantified by flow cytometry. MOI=multiplicity of infection. (**C**) Correlation of perturbed transcriptomes, (**D**) number of differentially expressed genes, (**E**) overexpression of intended targets, and (**F**) differential expression volcano plots for AAV overexpression of predicted age-promoting targets in human iPSC-derived cardiomyocytes. *q<0.05, Wald test with BH correction, n=4. (**G**) Overexpression of intended targets and (**H**) number of differentially expressed genes for AAV overexpression of predicted age-promoting targets in human primary cardiac fibroblasts. *q<0.05, Wald test with BH correction, n=4.

## Supplementary Tables

**Table S1.** Dataset composition of Genecorpus-175M.

**Table S2.** Gene set enrichment of cardiac fibroblast in silico predicted targets.

**Table S3.** Predicted impact on aging trajectory of in silico inhibition of gene targets in cardiac cell types.

**Table S4.** Differential gene expression by bulk RNA-sequencing in response to predicted pro-aging perturbations (AAV overexpression of indicated genes vs. GFP) in human iPSC-derived cardiomyocytes and human primary cardiac fibroblasts.

**Table S5.** Gene set enrichment of genes differentially expressed in response to all predicted pro-aging perturbations in human iPSC-derived cardiomyocytes.

**Table S6.** Gene set enrichment of genes differentially expressed in response to P4HA1 and RASGEF1B overexpression in both human iPSC-derived cardiomyocytes and human primary cardiac fibroblasts.

